# TERT prevents obesity-induced metabolic disorders by promoting adipose stem cell expansion and differentiation

**DOI:** 10.1101/2025.01.06.631346

**Authors:** Laura Braud, Manuel Bernabe, Julien Vernerey, Antonio M.A. Miranda, Andrea Dominguez, Dmitri Churikov, Manon Richaud, Liam Mc Allan, Christophe Lachaud, Jesus Gil, Will Scott, Vincent Géli

## Abstract

Obesity is linked to limited adipose tissue (AT) remodeling capacity, leading to hypertrophic adipocytes, senescence, and inflammation. We used a mouse model expressing *mTert* (p21^+/Tert^) from the Cdkn1a locus to investigate the role of mTERT in obesity-induced metabolic disorders. Conditional expression of mTERT reduces metabolic disorders associated with obesity. In AT, this is accompanied by a decrease in the number of senescent p21-positive cells, very short telomeres, and oxidative DNA damage. Single nucleus RNA-seq data reveal TERT expression attenuates senescence induced by HFD in particular in adipose stem and progenitor cells (ASPC). We show that ASPC expansion and differentiation are promoted in p21^+/Tert^ obese mice, thereby reducing metabolic disorders. We further report that mTERT remodels the landscape of macrophages in AT of obese mice. Strikingly, inactivation of mTERT catalytic activity in p21^+/Tert^ (p21^+/TertCi^) mice suppresses the promotion of adipocyte formation, but neither affects attenuation of senescence nor macrophage remodeling. These results highlight mTERT’s canonical and non-canonical functions in reducing obesity-associated metabolic disorders. Conditional expression of TERT thus appears as a potential therapeutic option for obesity.

## Introduction

Obesity and metabolic syndrome (MS) have increased dramatically in recent decades and is associated with increased risks of insulin-resistance, type 2 diabetes, cardiovascular diseases and cancer (Kompella & Vasquez, 2019). Obesity is characterized by white adipose tissue (WAT) expansion, a crucial organ in the pathology due to its fat storing capacity as well as its endocrine function, promoted by the secretion of several factors and hormones that influence whole body metabolism (Longo et al, 2019; Ronti et al, 2006). During obesity, numerous events participate in adipose tissue dysfunction, including adipocyte hypertrophy, impaired adipose stem and progenitor cell (ASPC) differentiation capacity, pro-inflammatory immune cell infiltration and fibrosis (Vishvanath & Gupta, 2019). ASPCs are found in the vascular niche of the adipose tissue (AT). They differentiate into mature adipocytes and confer a high degree of plasticity to the AT. There are distinct subtypes of ASPC in adipose tissue reflecting the complexity of ASPC in different adipose deposits (Liao et al, 2022). Obesity-induced progenitor dysfunction such as senescence and reduced adipogenic properties, lead to restricted capacity of AT remodeling and contribute to obesity associated metabolic disorders (Sakers et al, 2022).

The AT stromal vascular fraction (SVF) also includes adipose tissue macrophages (ATM), various classes of immune cells, fibroblasts, and endothelial cells, that play a crucial role in adipose tissue homeostasis (Tabula Muris Consortium et al, 2018; Jaitin et al, 2019; Merrick et al, 2019; Sárvári et al, 2021). ATMs are recruited by different factors secreted by the AT (McNelis & Olefsky, 2014). The recruited macrophages perform several functions both in health and disease, including elimination of dead adipocytes by phagocytosis, tissue homeostasis, insulin resistance, inflammation, and adipose tissue fibrosis (Chakarov et al, 2022). The ATM proportion in adipose tissue can rise from 10% to 40% in obese people (Weisberg et al, 2003). Of note, High Fat Diet (HFD)-fed mice massively produced lipid-associated macrophages (LAM) characterized by the expression of the lipid receptor Trem2. These macrophages, which are responsible for lipid absorption and metabolism, protect against adipocyte hypertrophy and inflammation (Sárvári et al, 2021).

These observations have led to a shift from the traditional view of adipose tissue as a passive energy storage reservoir to that of a complex and highly active metabolic and endocrine organ that needs to be finely regulated (Sakers et al, 2022).

Numerous studies in humans have shown that obesity and MS are associated with shortening of telomeres (Chen et al, 2014; Lakowa et al, 2015; Khosravaniardakani et al, 2022). This correlation between reduced telomere length and obesity is probably mediated by the oxidative stress and inflammation associated with obesity that can damage telomeres(Gavia-García et al, 2021; Gao et al, 2020; Palmer et al, 2019). Recent studies in mice demonstrated a direct link between obesity, senescence and telomeres. Indeed, specific inactivation of mTERT (the catalytic subunit of mouse telomerase) in mouse adipose progenitor cells results in premature telomere shortening and cellular senescence accompanied by adipocyte hypertrophy, inflammation, fibrosis and systemic insulin resistance (Gao et al, 2020). Besides, clearance of the p16^high^ senescent cells in obese mice has been shown to alleviate metabolic and adipose dysfunctions, implicating cellular senescence as a causal factor in obesity-related inflammation and metabolic disorders (Palmer et al, 2019). In a similar vein, it has been shown that in mice made obese by High Fat Diet (HFD), p21^Cdkn1a^ is highly expressed in visceral adipose tissue, with p21^high^ cells being mainly endothelial cells, macrophages, leukocytes and mesenchymal stem cells (Wang et al, 2021). Clearance of these p21high senescent cells in obese mice also alleviated adipose tissue dysfunctions and insulin-resistance (Wang et al, 2021, 20). Collectively, these studies revealed that senescent cells instigate obesity associated metabolic disorders (Kondoh & Hara, 2022). Of note, adipocytes from mice fed a high-fat diet show higher levels of oxidative DNA damage before the onset of alterations in adipocyte insulin sensitivity and glucose homeostasis (Kondoh & Hara, 2022). This is evidenced by an increase in cells with β-galactosidase activity, increased ATP content, the occurrence of DNA damage, and telomere attrition contributing to the activation of the Tp53-p21 pathway (Vergoni et al, 2016; Pini et al, 2021).

In humans, cellular senescence also increases in obese patients, as well as in patients suffering from type 2 diabetes or non-alcoholic fatty liver disease (Spinelli et al). Accumulation of senescent cells within adipose tissue is the main contributor to its dysfunctions (Conley et al, 2020; Palmer et al, 2015), and obesity-induced enlargement of adipocytes positively correlates with senescence (Palmer et al, 2015). Although elimination of cells expressing high levels of p21 and p16 reduced adipose tissue inflammation and improved insulin sensitivity in obese mice, much less is known about the consequences of telomere shortening and the role of canonical and non-canonical functions of TERT in adipose tissue function.

We recently generated p21^+/Tert^ transgenic mouse models in which telomerase reverse transcriptase (mTert), or its catalytically inactive form (mTertCi), is expressed from the p21Cdkn1a promoter (Lipskaia et al, 2024). We sought to enforce mTert expression specifically in p21-expressing cells, notably in pre-senescent cells including cells with dysfunctional telomeres. We have shown that this particular expression of mTert reduces accumulation of very short telomeres, oxidative damage, endothelial cells (ECs) senescence and senile emphysema in aged mice (Lipskaia et al, 2024). Unexpectedly, we found that both mTERT and mTERTCi expression significantly reduced p21 levels in the lungs of aged mice suggesting that a non-canonical function of mTert reduces p21 level (Lipskaia et al, 2024).

Here, we have taken advantage of these mouse models to study the canonical and non-canonical roles of mTERT and the impact of counteracting telomere attrition on obesity-induced metabolic disorders in mice fed with high fat diet (HFD). We show that mTERT expression under the control of the p21 promoter reduces the number of p21-expressing cells, the number of very short telomeres, inflammation, and senescence normally observed in HFD-fed control (p21^+/mCherry^) mice. This is accompanied by improved insulin sensitivity and glucose tolerance in obese mice. We show that mTERT stimulates ASPCs proliferation and differentiation into mature adipocytes in p21^+/Tert^ obese mice. This process contributes to the adipose tissue hyperplasia with increased number of adipocytes which has been shown to have a protective effect against obesity-associated metabolic disorders (Jang et al, 2016; Merrick et al, 2019). Moreover, we discovered that a non-canonical function of mTERT alters the landscape of macrophages in the AT of mice on HFD by reducing the populations of Trem2-positive lipid associated macrophages (LAM) while maintaining high levels of monocytes and tissue resident macrophages. Collectively, these findings reveal that canonical and non-canonical functions of TERT improves metabolic disorders mainly by inhibiting senescence signaling cascades, promoting ASPC expansion and differentiation into mature adipocytes, and attenuating the immune response.

## Results

### p21 promoter-dependent expression of mTERT prevents obesity-associated metabolic disorders in male obese mice

Because p21 is highly induced in mice fed with HFD (Wang *et al*, 2021), we evaluated the impact of the p21 promoter-driven expression of mTERT in obesity-induced metabolic disorders. The p21^+/Tert^ model in which the mCherry-2A-TERT cassette was inserted at the start codon of the *Cdkn1a*^p21^ gene has been described previously (Lipskaia *et al*, 2024). In addition, we generated here the p21^+/mCherry^ mouse as a control. These mice produce p21 protein from one allele and either mCherry alone or mTERT and mCherry from the other allele (Figure 1A).

**Figure 1.**
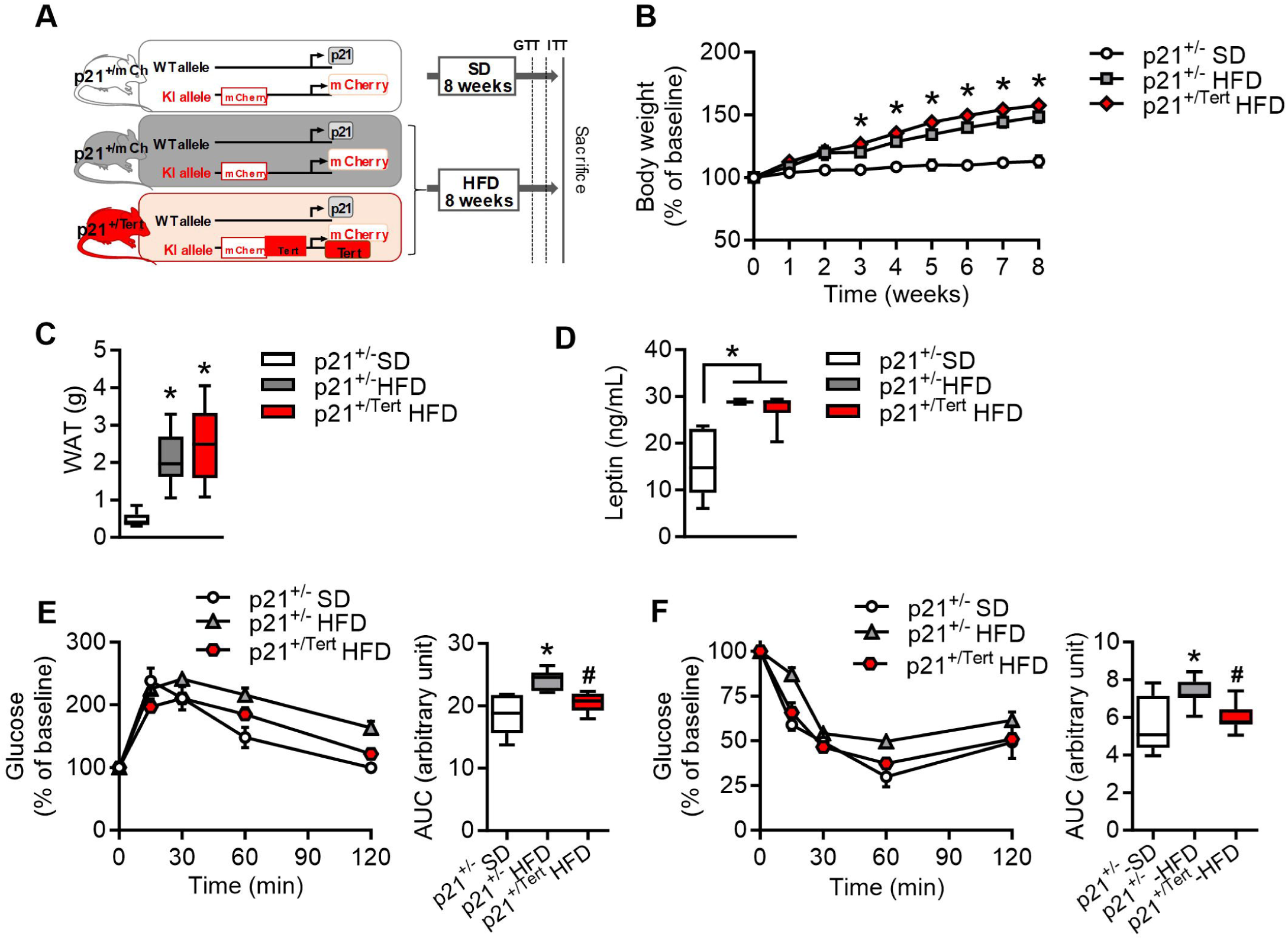
p21 promoter-driven mTert expression alleviates obesity-associated metabolic disorders. A. Schematic representation of mouse models B. Weight acquisition kinetics of the indicated HFD mice (n=7 p21^+/-^ SD, n=19 p21^+/-^ HFD and n=18 p21^+/Tert^ HFD mice) C. White adipose tissue (WAT) weight of HFD mice at sacrifice (n=6 p21^+/-^ SD, n=7 p21^+/-^ HFD and n=7 p21^+/Tert^ HFD mice) D. Leptin level in the blood plasma measured by Leptin ELISA (n=6 p21^+/-^ SD, n=5 p21^+/-^ HFD and n=6 p21^+/Tert^ HFD mice) E. Left, Glucose Tolerance Test (GTT) was performed by an intraperitoneal injection of glucose (1.5g/kg) and measurement of glycemia via tail clip (Caresens^®^ N, DinnoSanteTM) at different time points. Right, area under the curve (AUC) of the GTT (n=5 p21^+/-^ SD, n=8 p21^+/-^ HFD and n=7 p21^+/Tert^ HFD mice) F. Left, Insulin Tolerance Test (ITT) was performed by an intraperitoneal injection of insulin (0.3 UI/kg) and glucose was measured via tail clip (Caresens^®^ N, DinnoSanteTM) at different time points. Right, area under the curve (AUC) of the ITT (n=7 p21^+/-^ SD, n=9 p21^+/-^ HFD and n=11 p21^+/Tert^ HFD mice) All data are shown as mean ± SEM. *p<0.05 vs. p21^+/-^ SD mice (white bars), ^#^p<0.05 vs. p21^+/-^ HFD (grey bars). Student’s t test or one-way ANOVA with Fisher multiple comparison test.

We fed p21^+/mCherry^ male and female mice with standard chow diet (SD) as controls, and p21^+/mCherry^ and p21^+/Tert^ mice with HFD for 8 weeks (Figure 1A). Body weight gain was significantly increased in HFD groups compared to SD fed mice in both males and females (Figure 1B and Suppl Figure 1A). We did not observe any difference in body weight gain between p21^+/mCherry^ and p21^+/Tert^ obese mice in either males or females (Figure 1B and Suppl Figure 1A). Body weight gain was accompanied by increased accumulation of white adipose tissue (WAT) in all HFD mice, with no significant difference between groups (Figure 1C and Suppl Figure 1B). Accordingly, we also observed an increase in the plasma leptin level, the main adipokine secreted by AT, in all groups of HFD mice for both males and females with no difference between the genotypes (Figure 1d and Suppl Figure 1C).

In male mice, we observed that mTERT expression counteracted both whole-body glucose intolerance and insulin resistance induced by HFD, as evidenced by a significant decrease in glycemia compared to HFD-fed p21^+/mCherry^ mice Figure 1E-F). In female control mice (p21^+/mCherry^), HFD induced significant but very mild glucose intolerance and insulin resistance (Figure 1D-E). Therefore, we did not observe such improvement in p21^+/Tert^ obese female mice compared to p21^+/mCherry^ obese female mice (Figure 1D-E). Females are known to be more resistant against HFD induced metabolic disorders than male (Pettersson *et al*, 2012). Notably, we found that the level of p21 in the AT of control females is much lower than that in control males on SD and it does not increase on HFD (Figure 1F) that might explain the milder effect of mTERT expression on glucose metabolism in females. In all the following experiments, only the data for male mice are presented.

Thus, expression of mTERT under control of the p21 promoter leads to improved insulin resistance and reduced glucose intolerance in male mice fed a high-fat diet without any notable effect on the weight gain.

### Reduced accumulation of p21 positive cells in the SVF

As p21 has been shown to play a crucial role in adipose tissue dysfunction in HFD-fed mice (Wang *et al*, 2021), and because TERT expression reduces p21 levels (Lipskaia *et al*, 2024), we analyzed the number of mCherry-positive cells (reflecting p21 expression) in different cell types contained in the Stromal Vascular Fraction (SVF) namely pericytes, endothelial cells, pre-adipocytes, adipose stem and progenitor cells (ASPC), leukocytes, monocytes, dendritic cells (DC), macrophages that we defined at this stage as Macro CD11b^+^ CD11c^-^ and Macro CD11b^+^ CD11c^+^ (Murray, 2017). The different cell subtypes were identified by FACS analysis using cell surface markers and precise gating strategy (Suppl Figure 2). This enabled us to group the different cell subtypes and identify the mCherry-expressing cells in each group, based on the fluorescence emitted by the latter (Figure 2A). We found that the mCherry signal was significantly increased by HFD in all cell types with a greater increase in pre-adipocytes, ASPCs and macrophages (Figure 2A). We uncovered that the HFD-induced accumulation of mCherry-positive cells was reduced in p21^+/Tert^ obese mice in most of the SVF cell types (Figure 2A). Quantification of the fraction of mCherry positive cells in each cell type is shown in Figure 2B. Collectively, the results indicate that mTERT expression significantly reduces the fraction of p21-positive cells. We further analyzed p21 expression by qPCR in the SVF (Figure 2C). We found that global p21 expression level was higher in obese compared to lean mice (Figure 2C). mTERT expression significantly reduced p21 mRNA levels in mice fed HFD (Figure 2C).

**Figure 2.**
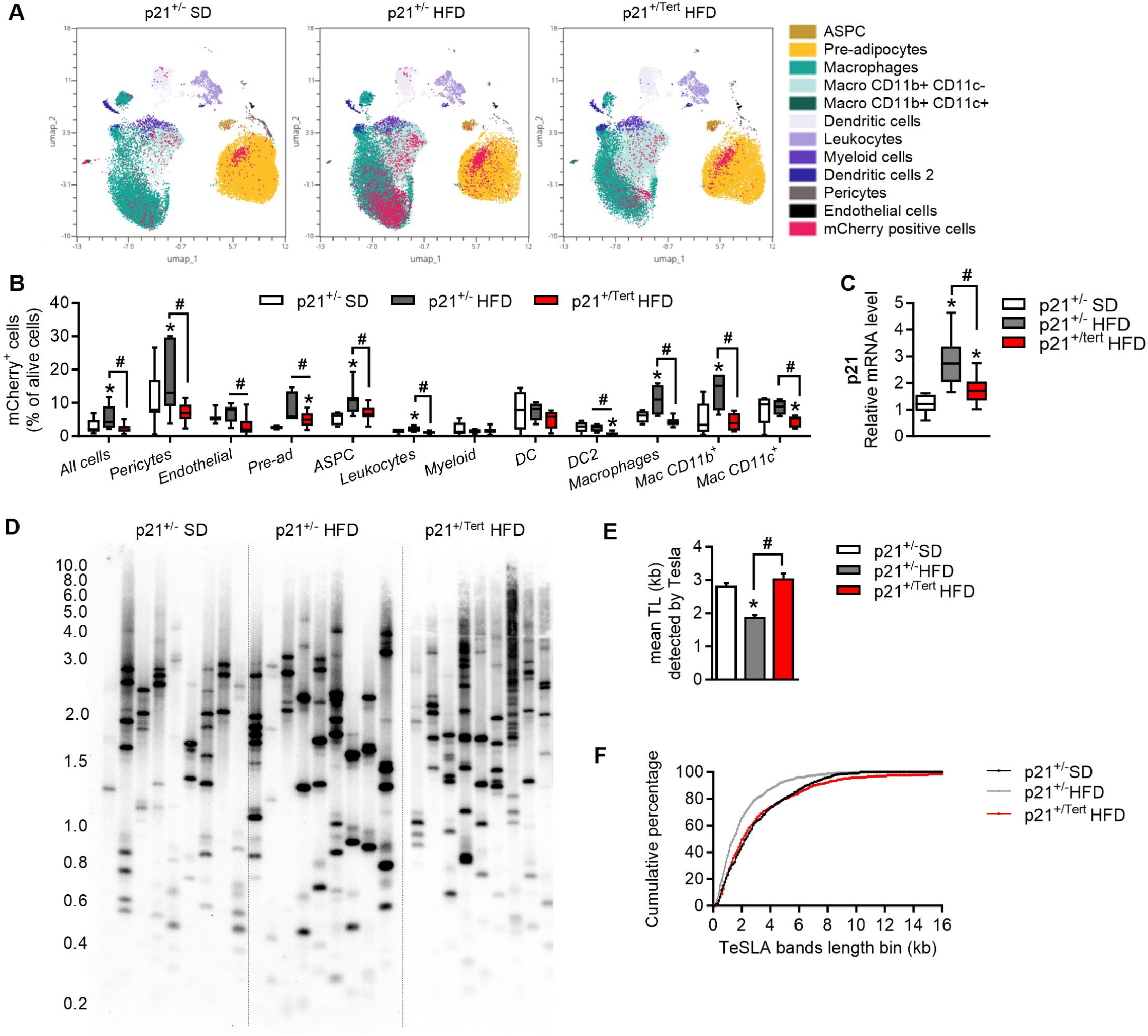
mTERT reduces the number of p21 positive cells in the stromal vascular fraction. A. UMAP clustering of SVF cells. Cell populations were distinguished by FACS using the markers indicated in Suppl Figure 2. mCherry fluorescence was measured by FACS and signal quantification in all cells is indicated. (n=6 mice per group) B. mCherry signal was measured by FACS. The percentage of mCherry positive cells is shown in the indicated populations of the SVF. (n=5-8 p21^+/-^ SD, n=4-7 p21^+/-^ HFD and n=10-17 p21^+/Tert^ HFD and mice) C. p21 mRNA levels was measured by RT-qPCR in stromal vascular fraction (SVF) (n=5 p21^+/-^ SD, n=6 p21^+/-^ HFD and n=6 p21^+/Tert^ HFD mice) D. Analysis of the short telomere fraction in the SVF cells isolated from the indicated mice by Telomere Shortest Length Assay (TeSLA). Genomic DNA was extracted from whole SVF; TeSLA was initiated with 50 ng of genomic DNA. In the final PCR step, 500 pg of ligation product was used per reaction, and 9 independent reactions were performed for each sample (each lane corresponds to an independent PCR reaction) to achieve > 100 amplified telomeres for quantification. The panel depicts one representative Southern blot for the indicated mice probed for the TTAGGG repeats. (n=3 per group) E. Mean telomere length of the telomeres detected by the Tesla method. (n=3 per group) F. Difference of cumulative number of short telomeres between SVF cells isolated from the indicated mice. (n=3 per group) All data are shown as mean ± SEM. *p<0.05 vs. p21^+/-^ SD mice (white bars), #p<0.05 vs. p21^+/-^ HFD (grey bars). Student’s t test or one-way ANOVA with Fisher multiple comparison test.

Finally, we analyzed the telomere length of the SVF cells in the control SD-fed mice and in the HFD-fed mice of the 2 genotypes. Because mice have long telomeres making it difficult to detect variations in telomere length by classical Southern blots, and because short telomeres instigate tissues dysfunction, we evaluated the presence of short telomeres using TeSLA (Lai *et al*, 2017) that allows detection of individual critically short telomeres in the bulk population of telomeres. As evident in the representative TeSLA (Figure 2D) and in the quantification graphs (Figure 2E-F), the obese control mice (p21^+/mcherry^) had on average more short telomeres than lean mice suggesting that the HFD-induced stress may damage telomeres. In contrast, the cumulative percentage of short telomeres were similar between the lean p21^+/mCherry^ and p21^+/Tert^ mice fed HFD suggesting that p21 promoter-dependent expression of mTERT heals the damaged telomeres (Figure 2D-E-F). Of note, we also detected HFD-induced telomere shortening and its attenuation by mTERT in total adipose tissue but it was less pronounced compared to that in SVF (Suppl Figure 3A-C). This may suggest that HFD affects telomere length primarily of the dividing cells in the SVF.

### mTERT downregulates genes involved in inflammation, angiogenesis, and extracellular matrix remodeling but upregulates oxidative phosphorylation genes

To assess the effects of p21 promoter-driven mTert expression on global transcription, we performed RNA-seq experiments on the SVF. We analyzed gene expression in the SVF of the p21^+/-^ and p21^+/Tert^ mice fed HFD and p21^+/-^ fed SD (Dataset 1). We first analyzed the genes which expression changes between SD- and HFD-fed p21^+/-^ mice. The volcano plot depicts the top 20 up- and down-regulated genes among the 654 and 236 genes that are significantly up- and down-regulated, respectively, reflecting the high plasticity of the AT (Maniyadath *et al*, 2023) (Suppl Figure 4A, left panel). The Gene Ontology (GO) analysis identified 54 biological processes (with adjusted p values ranging from 10^-22^ to 0,0025) (Suppl Figure 4A, right panel; Dataset 1 GO). In Figure 3A the 10 most significant biological processes with corresponding top 10 differentially expressed genes are shown. Deregulated genes are involved in different processes previously associated with obesity such as extracellular structure organization, remodeling of the immune system, and angiogenesis. We found that all these genes are up-regulated in the SVF of p21^+/-^ mice fed HFD. Remarkably, expression of mTERT attenuates the upregulation of all these genes induced by HFD (Figure 3A). In particular, we found that a number of events related to immune system activation in the adipose tissue of HDF-fed mice are attenuated in p21^+/Tert^ mice.

**Figure 3.**
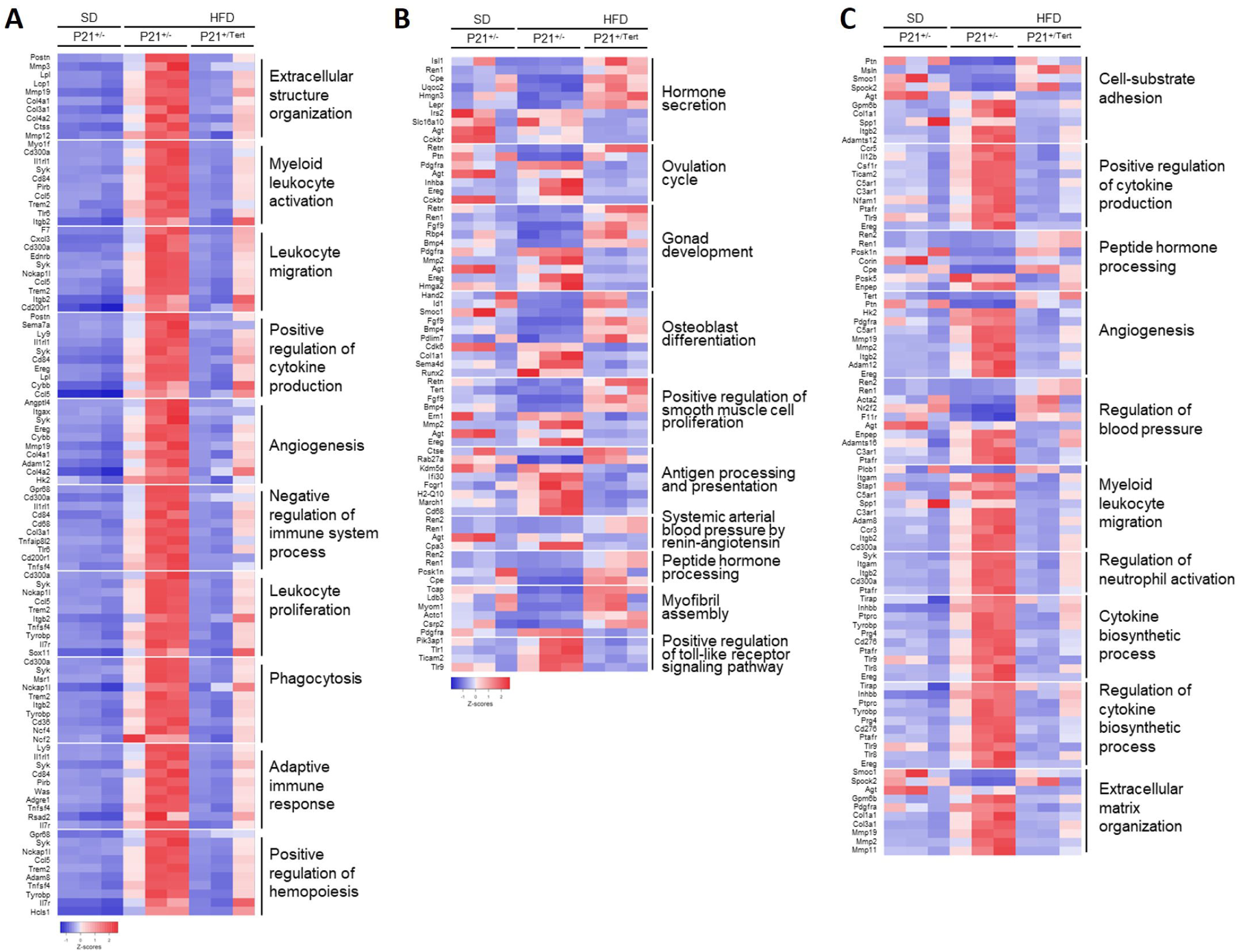
The adipose tissue SVF is remodeled in p21^+/Tert^ HFD mice. A. Heatmap showing the 10 most significant biological processes for which the most differentially regulated genes are represented between p21^+/-^ SD vs p21^+/-^ HFD B. Heatmap showing the 10 most significant biological processes for which the most differentially regulated genes are represented between p21^+/-^ HFD vs p21^+/Tert^ HFD C. Heatmap showing the 10 most significant biological processes for which the most differentially regulated genes are represented between p21^+/-^ HFD vs p21^+/TertCi^ HFD

We then compared SVF bulk transcriptomes between p21^+/-^ and p21^+/Tert^ HFD fed mice. Volcano plots (Suppl Figure 4B) show the top 20 up- and down-regulated genes in the SVF of HFD-fed p21^+/Tert^ mice compared with HFD-fed p21^+/-^ mice and the corresponding GO terms. As above, we show the 10 most significant biological processes (Figure 3B, Dataset 1). For each of the 2 comparisons (p21^+/-^ versus p21^+/Tert^), we again represented the most differentially expressed genes for each biological process (Figure 3B, Dataset 1). The GO analysis revealed that expression of mTERT differentially regulates genes involved in processes such as hormone secretion, osteoblast differentiation, and regulation of smooth muscle cell proliferation (Figure 3B). Of note, the genes participating in the osteoblast differentiation are either down- (Runx2) or up-regulated (Bmp4) in the SVF of p21^+/Tert^ mice fed with HFD (Figure 3B, Dataset 1). Interestingly, we found upregulation of the receptor of leptin (LepR), known to stimulate adipogenesis, in the pre-adipocytes (Palhinha *et al*, 2019) of the p21^+/Tert^ HFD-fed mice. As already shown in Figure 3a, we find that upregulation of genes involved in positive regulation of angiogenesis and interactions between immune cells was attenuated by mTERT expression (Figure 3C). Interestingly, we found that mTERT expression resulted in the up-regulation of genes involved in oxidative phosphorylation and proton motive force-driven ATP synthesis such as Cox6c, Uqcr10, Cox6b2, Uqcrh, Atp5j2, Ndufa12, NduFab1, and Fmo3 (Dataset 1). Overall, mTERT expression in HFD mice downregulates a number of processes associated with inflammation, angiogenesis, and extracellular matrix organization and appear to promote healthy energy metabolism.

### mTERT expression alleviates adipose progenitor senescence induced by HFD

To assess the effect of mTERT expression controlled by the p21 promoter in specific AT cell types, we performed single-nucleus RNA sequencing (snRNA-seq) on WAT from p21^+/mCherry^ mice fed with SD and p21^+/mCherry^ and p21^+/Tert^ fed with HFD for 8 weeks. After quality filtering, we generated dataset for p21^+/-mCherry^ SD (n=2), p21^+/mCherry^ HFD (n=2), and p21^+/Tert^ HFD (n=2). We integrated the datasets and performed dimensionality reduction and unsupervised cell clustering to identify distinct cell types based on unique and shared gene expression patterns (Emont *et al*, 2022; Sárvári *et al*, 2021) (Figure 4A). The number of cells for each cell type and for each condition is indicated in the Dataset 2. For all mice, we recovered nuclei from the main cell types reported to be present in the WAT (Emont *et al*, 2022; Wang *et al*, 2021). We analyzed p21 expression in the different AT cell types in p21^+/mCherry^ and p21^+/Tert^ mice fed with HFD (Figure 4B). In HFD-fed control mice (p21^+/mCherry^), p21 expression was mainly induced in ASPCs and macrophages (Figure 4B). In agreement with our observations in SVF (Figure 2), the fraction of p21-positive cells was markedly reduced in all AT cell types of p21^+/Tert^ HFD mice (Figure 4B).

**Figure 4.**
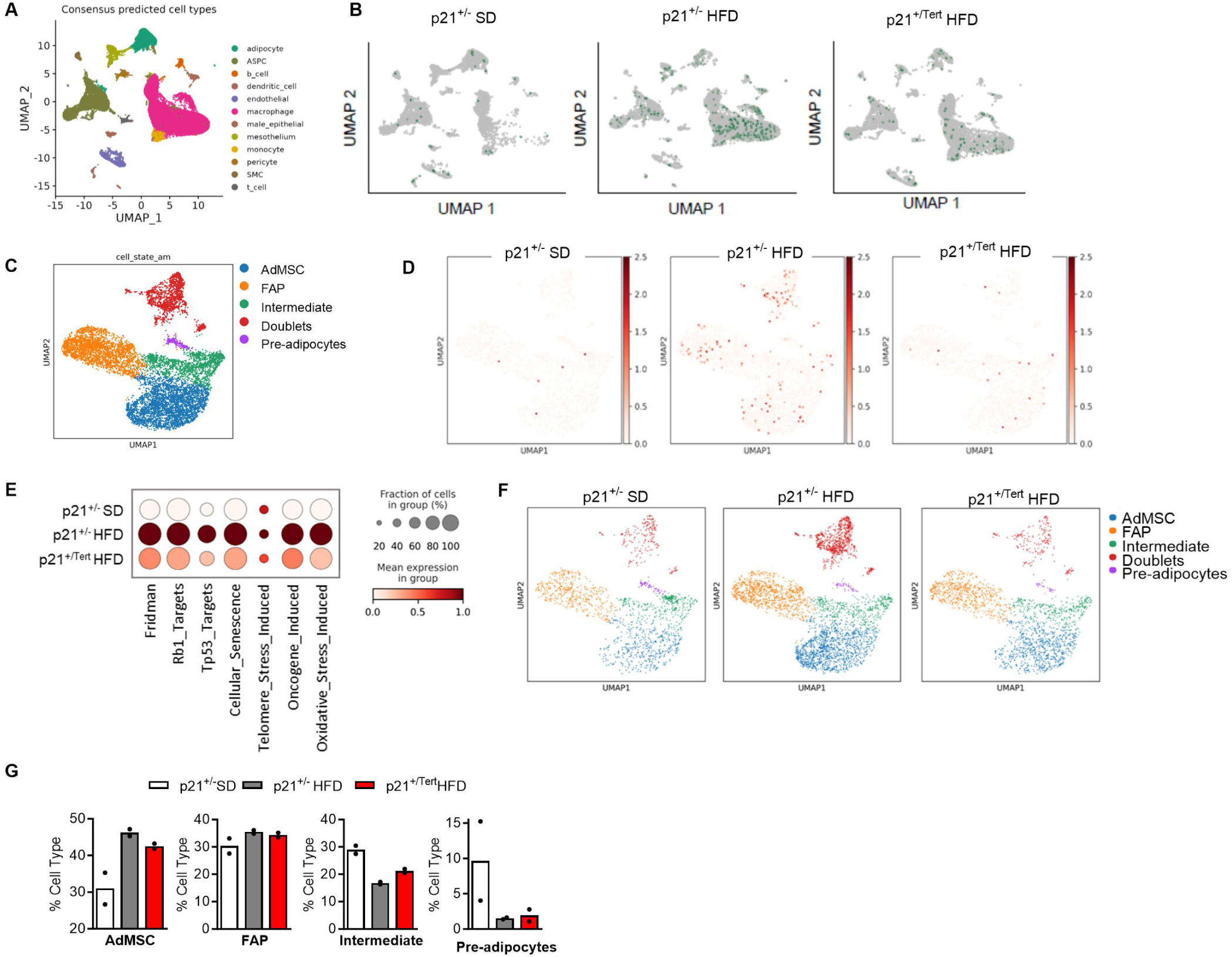
mTERT expression alleviates cellular senescence in ASPC. A. snRNAseq UMAP of the cell populations in all cells B. snRNAseq UMAP of the p21 expression in the different cell populations in p21^+/-^ SD, p21^+/-^ HFD and p21^+/Tert^ HFD mice C. Subclustering of ASPC from snRNAseq data D. p21 expression in different subtypes of adipocytes progenitors in p21^+/-^ SD, p21^+/-^ HFD and p21^+/Tert^ HFD mice E. Scoring of senescence pathways in the different group of mice in the progenitors F. Representative ASPC subtypes from the WAT of p21^+/-^ SD, p21^+/-^ HFD and p21^+/Tert^ HFD mice G. Mean percentage of ASPC subtypes per sample (excluding doublets)

Since adipose stem and progenitor cells are critical in adipose tissue homeostasis, we performed a detailed analysis of the ASPC population. ASPCs were subclustered using unsupervised clustering and selected markers (Suppl Figure 5) resulting in 4 subtypes (Adipose Mesenchymal Stem Cells-AdMSC, Fibro-adipogenic progenitors-FAP, Intermediate progenitors-Inter and Pre-adipocytes-Pread) (Figure 4C) according to Cannavino and Gupta (Cannavino & Gupta, 2023). We found that *Cdkn1a* (the transcript encoding p21) mRNA levels were elevated specifically in FAP and AdMSC, of p21^+/mCherry^ mice fed HFD, but this increase was diminished in p21^+/Tert^ HFD-fed mice (Figure 4D). In p21^+/mCherry^ HFD-fed mice, we also found that *Cdkn1a* was upregulated in other cell types in particular in adipocytes and myeloid cells mice and that this increase was also diminished in p21^+/Tert^ HFD-fed mice (Suppl Figure 6A-B).

The fact that the number of p21-positive cells was markedly reduced in ASPCs of p21^+/Tert^ HFD mice prompted us to assess senescence signatures in the different mice fed with HFD. We revealed increased expression of the senescence-associated genes, notably targets of Tp53 and Rb in the ASPCs of p21^+/mCherry^ mice exposed to HFD. Strikingly, up-regulation of the senescence genes was strongly decreased in ASPCs from p21^+/Tert^ mice (Figure 4E). In addition, the increase in senescence associated with HFD is also diminished in other cell types derived from the fat of p21^+/Tert^ mice such as adipocytes and myeloid cells (Suppl Figure 6C-E). Overall, these results suggest that p21-promoter dependent expression of *mTert* alleviates the accumulation of senescent cells in ASPCs and other cell types in obese mice.

We then sought to identify differentially expressed genes (DEGs) between p21^+/mCherry^ and p21^+/Tert^ ASPCs from mice fed HFD. To perform a more robust assessment of DEGs, for each gene within each condition we sum the data of the 2 single nucleus RNA-seq experiments (that gave very similar expression levels, Dataset 3) and used a statistical method (DESeq2). The volcano plots in Suppl Figure 7 show the 20 most up- and down-regulated genes between ASPCs from p21^+/mCherry^ and p21^+/Tert^ mice. These genes include (Fat3, Lrrc4, Ftl1, Fth1, Mmp12, Apoe, Fgf13, Cd84, Ctsl, Lgals3, Apobec1, Strbp, Fgfr2, Igf2r) (See also Dataset 3). Interestingly, it was very recently reported that the specific inactivation of Fth1 in adipose tissue is accompanied by a decrease in mitochondrial ROS levels, increased insulin sensitivity and glucose tolerance (Lu *et al*, 2024) (see discussion). On the other hand, we found that genes described as negative regulators of the Wnt /β-catenin signaling pathway in the GO database were overexpressed in p21^+/Tert^ ASPC (Dataset 3). This is illustrated by the up-regulation of Dact1, Wnt5b, Tnem64 that have been reported to promote adipogenesis by counteracting activation of the Wnt pathway (Lagathu *et al*, 2009; van Tienen *et al*, 2009; Jeong *et al*, 2015).

Finally, we looked at the ASPC subtypes proportion. We found that the percentage of AdMSC (less differentiated) was slightly reduced in the WAT of p21^+/Tert^ obese mice compared to the obese p21^+/mCherry^ mice while the percentage of intermediate progenitors was slightly higher (Figure 4F-G) suggesting that TERT promotes the differentiation of ASPCs.

### mTERT inhibits pre-adipocytes senescence in vitro and improves their differentiation

We aimed at consolidating the above results by culturing pre-adipocytes isolated from the 3 different groups of mice. Briefly, WAT was collected from SD and HFD p21^+/mCherry^ mice and from HFD p21^+/Tert^ mice to collect the stromal vascular fraction and the cells were plated in 6 well plates. At confluence, the differentiation of pre-adipocytes into mature adipocytes was stimulated by medium containing insulin. We observed that differentiation was reduced in p21^+/mCherry^ obese mice compared to pre-adipocytes isolated from SD-fed p21^+/mCherry^ mice (Figure 5A). This differentiation defect was attenuated in preadipocytes from p21^+/Tert^ obese mice (Figure 5A). In line with these results, expression of the adipogenic genes (adiponectin, Glut4 and Pparγ) was significantly enhanced in p21^+/Tert^ pre-adipocytes compared to p21^+/mCherry^ pre-adipocytes (Figure 5B), confirming improved differentiation of preadipocytes into mature adipocytes in p21^+/Tert^ obese mice compared with obese p21^+/mCherry^ mice. We then performed senescence-associated β-galactosidase (SA-β-Gal) staining of pre-adipocytes to assess senescence induction. In agreement with previous studies (Pini *et al*, 2021; Narasimhan *et al*, 2022), we observed a greater number of SA-β-galactosidase-positive cells in preadipocytes from obese p21^+/mCherry^ mice compared with preadipocytes from p21^+/mCherry^ mice fed in SD, reflecting an increase in senescence caused by the HFD (Schafer *et al*, 2016) (Figure 5C-D). In contrast, preadipocytes derived from p21^+/Tert^ obese mice did not show elevated levels of senescence (Figure 5C-D). In line with our previous results, p21^+/mCherry^ HFD preadipocytes showed higher p21 expression than those from SD-fed mice (Figure 5E). We again observed a lower percentage of preadipocytes positive for p21 in p21^+/Tert^ mice when compared to that in p21^+/mCherry^ mice (Figure 5E). We wondered whether the senescence observed in the preadipocytes of obese p21^+/mcherry^ mice correlated with telomere shortening in these mice. We thus analyzed short telomeres by TeSLA (Lai *et al*, 2017) in the cultured preadipocytes from SD and HFD mice of the 3 genotypes. We observed that preadipocytes from obese p21^+/mcherry^ mice had shorter telomeres than preadipocytes from lean mice (Figure 5F-H). Similar to what we observed in SVF (Figure 2D), HFD-induced telomere shortening was counteracted by mTERT expression in preadipocytes from obese p21^+/Tert^ mice (Figure 5F-H). We infer from the TeSLA results that p21^+/mcherry^ pre-adipocytes senescence might be driven by increased telomeric damage (Jacobs & de Lange, 2004).

**Figure 5.**
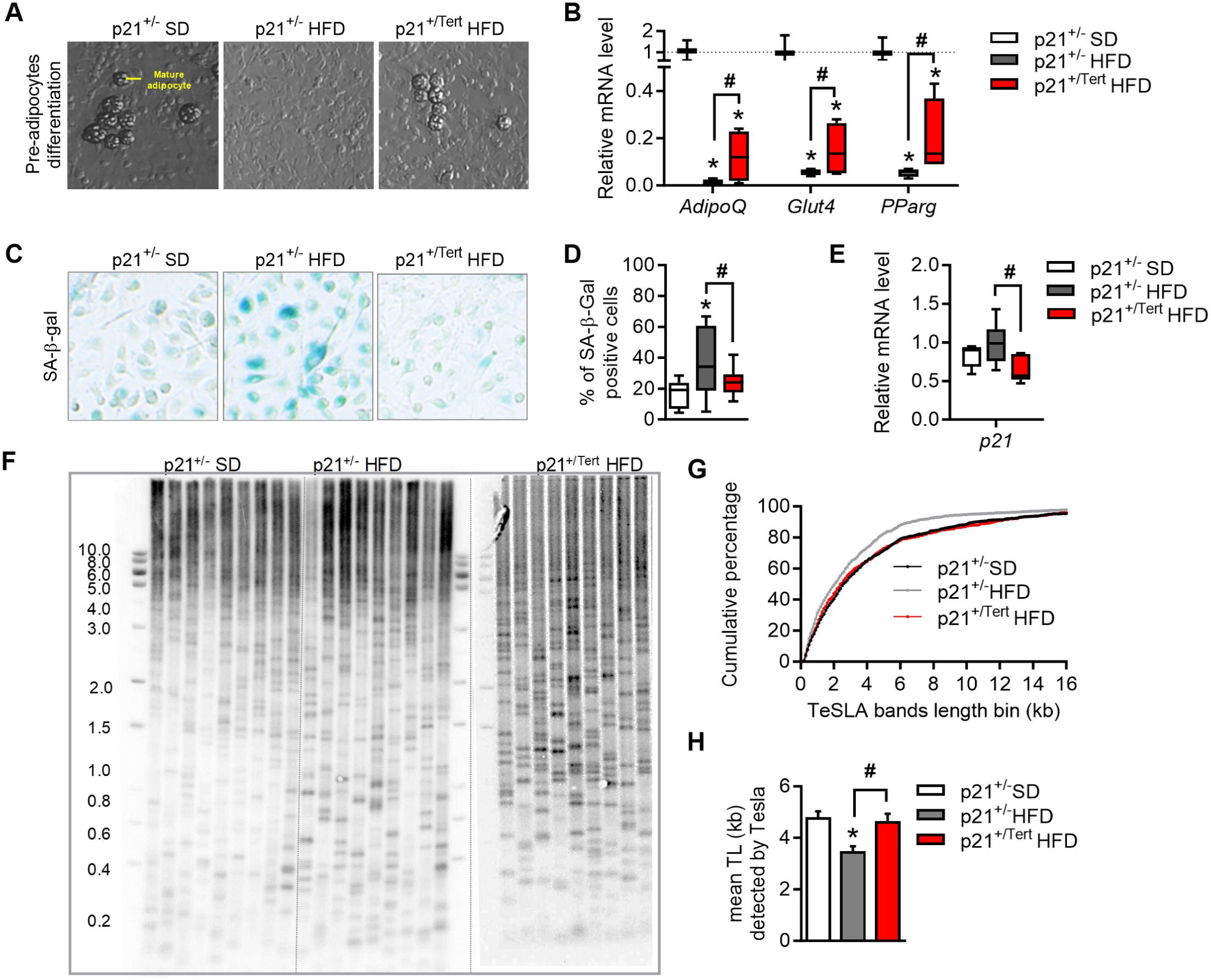
Pre-adipocytes isolated from SVF of HFD p21^+/Tert^ mice show reduced senescence and enhanced differentiation. A. Representative photographs showing adipocytes formation from pre-adipocytes in cell culture B. Expression of the genes involved in lipogenesis was measured by RT-qPCR in pre-adipocytes cell cultures established from WAT of the p21^+/-^ SD, p21^+/-^ HFD and p21^+/Tert^ HFD mice (n=3 p21^+/-^ SD, n=4 p21^+/-^ HFD and n=4 p21^+/Tert^ HFD mice) C. SA-β-gal staining of cultured pre-adipocytes from the WAT of the p21^+/-^ SD, p21^+/-^ HFD and p21^+/Tert^ HFD mice (n=3 p21^+/-^ SD, n=4 p21^+/-^ HFD and n=4 p21^+/Tert^ HFD mice) D. Quantification of SA-β-gal positive cells in the cultures established from WAT of the indicated groups of mice (n=3 p21^+/-^ SD, n=4 p21^+/-^ HFD and n=4 p21^+/Tert^ HFD mice) E. p21 mRNA expression was measured by RT-qPCR in pre-adipocytes cell cultures established from WAT of the p21^+/-^ SD, p21^+/-^ HFD and p21^+/Tert^ HFD mice (n=5 p21^+/-^ SD, n=7 p21^+/-^ HFD and n=7 p21^+/Tert^ HFD mice) F. The shortest telomere burden in pre-adipocytes analyzed by TeSLA (See Figure 2e for details) (n=3 per group) G. Difference between the cumulative numbers of short telomeres for the cultured pre-adipocytes isolated from the SVF of the indicated mice. Three independent cultures were analyzed for the indicated mice. (n=3 per group) H. Mean telomere length of the telomeres detected by the Tesla method in the indicated groups of mice (n=3 per group) All data are shown as the mean ± SEM. *p<0.05 vs. p21^+/-^ SD mice (white bars), #p<0.05 vs. p21^+/-^ HFD (grey bars). Student’s t test or one-way ANOVA with Fisher multiple comparison test.

### Formation of mature adipocytes is improved in p21^+/Tert^ obese mice

To further confirm the effect of TERT on adipocyte formation, we performed a histological analysis of the adipose tissue of the HFD mice. We first noticed that, despite no difference in WAT weight between obese mice (Figure 1C), the adipocyte size in the WAT of obese p21^+/Tert^ mice was reduced, with smaller and denser adipocytes compared to obese control mice (Figure 6A-D). In addition, we found that expression (analyzed by qPCR) of a number of adipocyte markers [adiponectin (AdipoQ), Glucose transporter 4 (Glut4), peroxisome proliferator-activated receptor gamma (PPARγ), fatty acid synthase (Fasn), sterol regulatory element-binding transcription factor 1 (Srebp1c), and Insulin receptor 1 (Irs1)] was down-regulated in p21^+/mCherry^ fed with HFD, whereas it was partially maintained in p21^+/Tert^ HFD mice (Figure 6E). FACS analysis of the WAT revealed an accumulation of ASPC in the SVF of p21^+/mCherry^ obese mice while the number of ASPC in obese p21^+/Tert^ mice was at the same level as in SD-fed control mice (Figure 6F). Conversely, the number of preadipocytes was similar in the p21^+/mCherry^ and p21^+/Tert^ obese mice (Figure 6G). Collectively, these results suggest that mature adipocyte formation is improved in p21^+/Tert^ obese mice.

**Figure 6.**
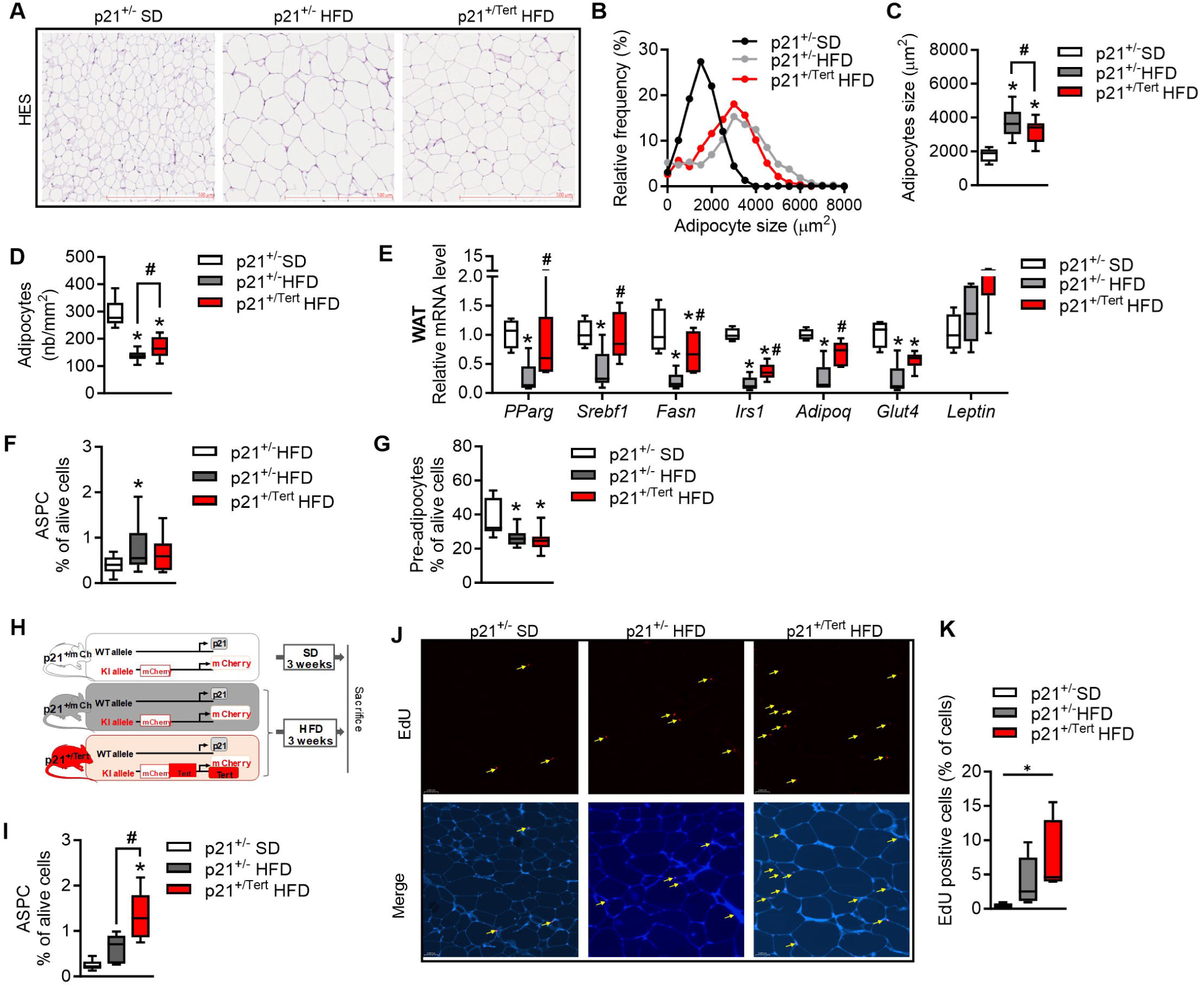
mTERT improves formation of adipocytes in obese mice. A. Hematoxylin and eosin (H&E) staining of visceral adipose tissue (WAT) from p21^+/-^ SD, p21^+/-^ HFD and p21^+/Tert^ HFD mice. B. Relative frequency of adipocyte sizes in the indicated groups of mice was plotted using GraphPad (n=5 p21^+/-^ SD, n=13 p21^+/-^ HFD and n=15 p21^+/Tert^ HFD mice) C. Adipocytes size in the images of H&E-stained WAT was quantified using AdipoSoft ImageJ software (n=5 p21^+/-^ SD, n=13 p21^+/-^ HFD and n=15 p21^+/Tert^ HFD mice) D. Adipocyte cell density in the images of H&E-stained WAT was quantified using AdipoSoft ImageJ software (n=5 p21^+/-^ SD, n=13 p21^+/-^ HFD and n=15 p21^+/Tert^ HFD mice) E. Expression of the genes involved in lipogenesis, lipolysis and fatty acid oxidation was measured by RT-qPCR in WAT of the p21^+/-^ SD, p21^+/-^ HFD and p21^+/Tert^ HFD mice (n=5 p21^+/-^ SD, n=6 p21^+/-^ HFD and n=6 p21^+/Tert^ HFD mice) F. Relative percentage of Adipose Stem and Progenitors Cells (ASPC) in the WAT of the p21^+/-^ SD, p21^+/-^ HFD and p21^+/Tert^ HFD mice analyzed by FACS (n=13 p21^+/-^ SD, n=11 p21^+/-^ HFD and n=15 p21^+/Tert^ HFD mice) G. Relative percentage of pre-adipocytes in the WAT of the p21^+/-^ SD, p21^+/-^ HFD and p21^+/Tert^ HFD mice analyzed by FACS (n=13 p21^+/-^ SD, n=11 p21^+/-^ HFD and n=15 p21^+/Tert^ HFD mice) H. Scheme of new protocol for early time HFD (3 weeks HFD) I. Relative percentage of Adipose Stem and Progenitors Cells (ASPC) in the WAT of the p21^+/-^ SD, p21^+/-^ HFD and p21^+/Tert^ HFD mice analyzed by FACS (n=6 p21^+/-^ SD, n=6 p21^+/-^ HFD and n=7 p21^+/Tert^ HFD mice) J. EdU staining into APSC from male WAT from p21^+/-^ SD, p21^+/-^ HFD and p21^+/Tert^ HFD mice after 48 hours pulses of EdU (n=4 p21^+/-^ SD, n=5 p21^+/-^ HFD and n=4 p21^+/Tert^ HFD mice) K. Quantification of EdU positive cells (n=4 p21^+/-^ SD, n=5 p21^+/-^ HFD and n=4 p21^+/Tert^ HFD mice) All data are shown as the mean ± SEM. *p<0.05 vs. p21^+/-^ SD mice (white bars), ^#^p<0.05 vs. p21^+/-^ HFD (grey bars). Student’s t-test or one-way ANOVA with Fisher multiple comparison test.

To determine whether mTERT stimulates ASPC proliferation in the WAT of p21^+/Tert^ mice fed with HFD, we fed the mice with HFD over a shorter period (3 weeks instead of 8) (Figure 6H). Indeed, ASPC proliferation has been described as appearing at the beginning of the HFD (Jeffery *et al*, 2015). After 3 weeks of HFD, FACS analysis reveal a greater accumulation of ASPC in the SVF of p21^+/Tert^ obese mice compared with p21^+/mcherry^ obese mice (Figure 6J). To confirm this result, mice were injected intraperitoneally with clickable EdU after 3 weeks of HFD and 48 hours before sacrifice. EdU was stained by click-it reaction on paraffin sections and we analyzed EdU-positive cells in the AT of the mice. We found that p21^+/mCherry^ and p21^+/Tert^ of obese mice showed an increase in the number of EdU-positive cells compared with SD mice (Figure 6J-K). However, we found a 2-fold increase in EdU-positive cells in p21^+/Tert^ obese mice, confirming that mTERT stimulates ASPC proliferation *in vivo* (Figure 6J-K).

### mTERT expression improves mitochondrial function and reduces oxidative stress

Telomere dysfunction in mice has been associated with impaired mitochondrial function and increased reactive oxygen species (Sahin *et al*, 2011; Amano *et al*, 2019). Moreover, in cell models, TERT has been shown to localized in mitochondria upon oxidative stress and to decrease ROS generation (Indran *et al*, 2011; Saretzki, 2014). These studies combined to the above results prompted us to assess the impact of p21 promoter-driven expression of mTERT on the bioenergetics of the cultured pre-adipocytes. Using the Seahorse analyzer, which measures the rate of oxygen consumption in the cell culture medium, reflecting the mitochondrial function of the cells, we found that preadipocytes isolated from p21^+/mCherry^ obese mice showed a slight decrease in basal and ATP-related respiration compared with preadipocytes isolated from SD p21^+/mCherry^ mice (Figure 7A-B) indicating that HFD treatment induces mitochondrial dysfunction in preadipocytes. Strikingly, we observed that p21^+/Tert^ pre-adipocytes displayed similar respiratory rates than pre-adipocytes isolated from p21^+/mCherry^ mice fed in SD (Figure 7A-B) suggesting that that conditional expression of mTERT preserves mitochondrial function in mice fed with HFD. Of note, we also observed that glycolysis was also preserved in p21^+/Tert^ pre-adipocytes compared to p21^+/mCherry^ pre-adipocytes from obese mice (Figure 7C).

**Figure 7.**
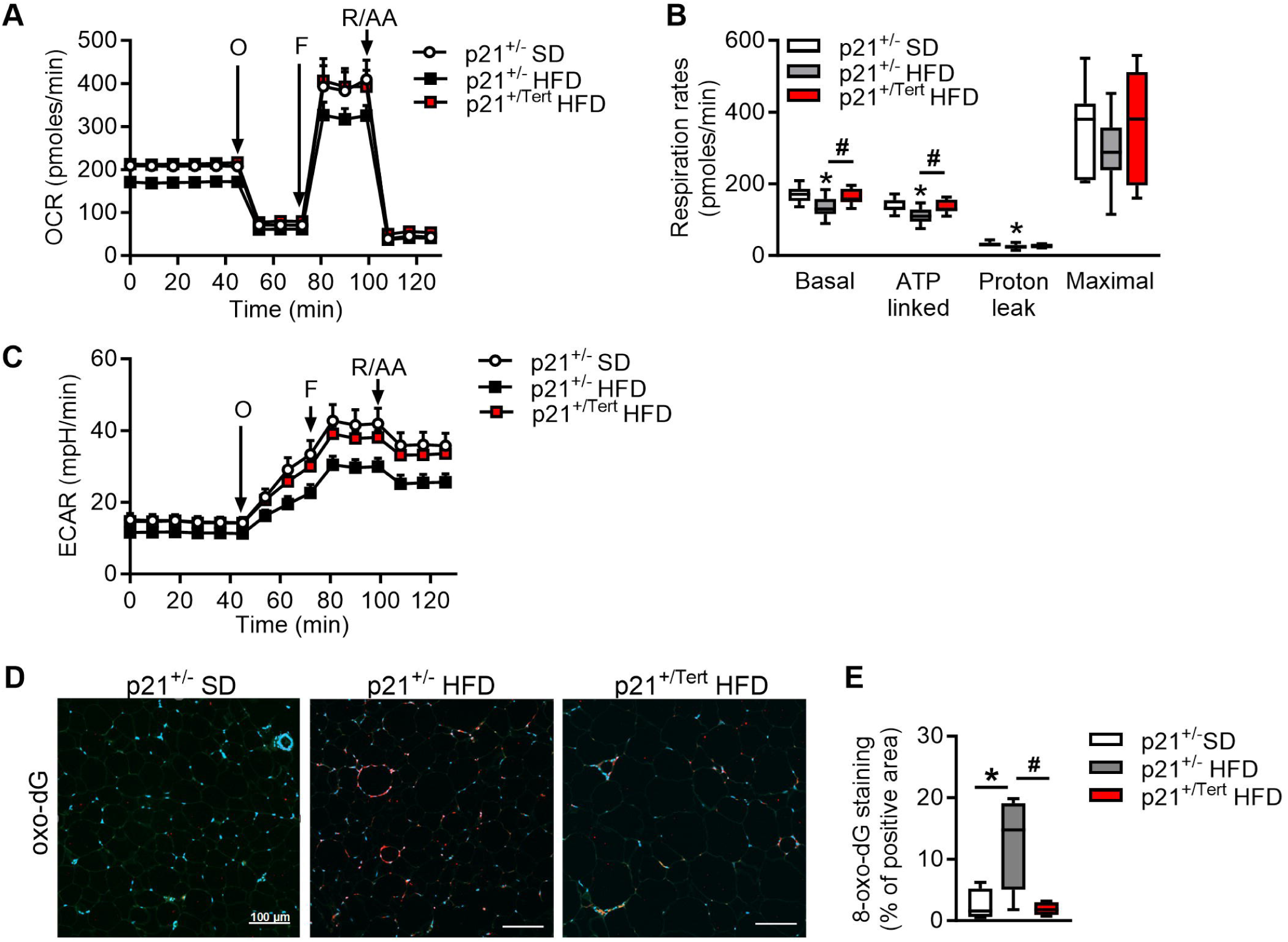
mTert expression reduces oxidative stress and mitochondrial dysfunction induced by obesity. A. Oxygen consumption rate analyzed by Seahorse in ASPC cell cultures established from WAT of the p21^+/-^ SD, p21^+/-^ HFD and p21^+/Tert^ HFD mice (n=3 p21^+/-^ SD, n=4 p21^+/-^ HFD and n=4 p21^+/Tert^ HFD mice in duplicate) B. Respiratory rates calculated based on oxygen consumption data (n=3 p21^+/-^ SD, n=4 p21^+/-^ HFD and n=4 p21^+/Tert^ HFD mice in duplicate) C. Extracellular acidification rate (ECAR), an index of glycolysis (n=3 p21^+/-^ SD, n=4 p21^+/-^ HFD and n=4 p21^+/Tert^ HFD mice in duplicate) D. 8-Oxo-désoxyguanosine (oxo-dG) staining of visceral adipose tissue (WAT) from p21^+/mcherry^ SD, p21^+/mCherry^ HFD and p21^+/Tert^ HFD mice. (n=3 p21^+/-^ SD, n=4 p21^+/-^ HFD and n=4 p21^+/Tert^ HFD mice) E. Quantification of the Oxo-dG positive cell (n=3 p21^+/-^ SD, n=4 p21^+/-^ HFD and n=4 p21^+/Tert^ HFD mice) All data are shown as the mean ± SEM. *p<0.05 vs. p21^+/-^ SD mice (white bars), #p<0.05 vs. p21^+/-^ HFD (grey bars). Student’s t test or one-way ANOVA with Fisher multiple comparison test.

We have previously shown that the overall level of oxidative damage was reduced in the lungs of aged p21^+/Tert^ mice (Lipskaia *et al*, 2024). We therefore wondered whether this was also the case in the AT of p21^+/Tert^ mice fed with HFD. We analyzed the overall level of oxidative DNA damage by measuring 8-oxo-dG levels in the AT of p21+/^mCherry^ and p21^+/Tert^ mice fed with HFD. We found that 8-oxo-dG levels were increased in the AT of HFD-fed p21^+/mCherry^ mice, but that this increase in oxidative damage was abolished in obese p21^+/Tert^ mice (Figure 7D-E).

In summary, expression of TERT under the control of the p21-promoter strongly reduces DNA oxidative damage and improves mitochondrial function and glycolysis in preadipocytes cultured *in vitro* harvested from the AT of obese mice.

### Inactivation of TERT catalytic activity only partially alleviates metabolic disorders in obese mice but still attenuates p21 expression and senescence

Different studies based on conditional TERT overexpression have revealed TERT functions not associated with its telomere elongation activity (Saretzki, 2014; Sarin *et al*, 2005). In order to determine whether TERT’s catalytic activity is necessary for the different phenotypes we have described above, we used the p21^+/TertCi^ model that produces a catalytically inactive telomerase (TERT^Ci^) instead of TERT (Lipskaia *et al*, 2024). Consistent with the lack of mTERT activity, we found that the cumulative percentage of short telomeres was similar in the SVF fraction from p21^+/mCherry^ and p21^+/TertCi^ mice fed with HFD (Figure 8A). We found that mTERT^Ci^ expression had a mild effect on glucose intolerance but did not improve insulin resistance induced by HFD indicating that the ability of mTERT to synthesize telomeres is important to counteract the metabolic defects induced by HFD (Figure 8B-C). This is also reflected by the lack of difference between p21^+/mCherry^ and p21^+/TertCi^ obese mice on adipocyte size, density within the AT, cell proliferation, and adipogenic gene expression (Figure 8D-E-F-G).

**Figure 8.**
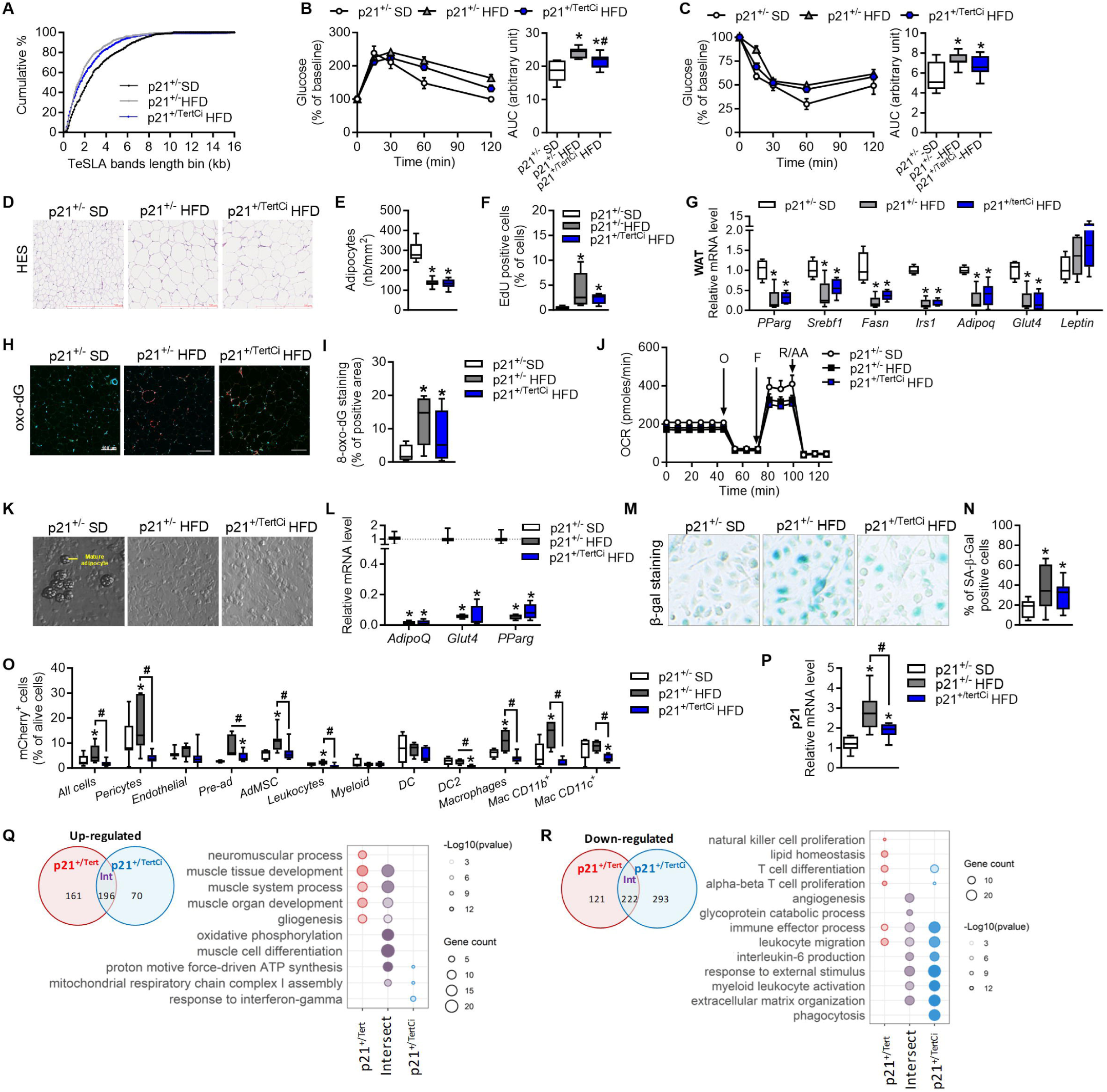
Inactivation of TERT catalytic activity only partially alleviates metabolic disorders in obese mice but still attenuates p21 expression. A. Difference between the cumulative numbers of short telomeres for the cultured pre-adipocytes isolated from the SVF of the indicated mice. Three independent cultures were analyzed for the indicated mice. (n=3 per group) B. Left, Glucose Tolerance Test (GTT) was performed by an intraperitoneal injection of glucose (1.5g/kg) and measurement of glycemia via tail clip (Caresens^®^ N, DinnoSanteTM) at different time points. Right, area under the curve (AUC) of the GTT (n=5 p21^+/-^ SD, n=8 p21^+/-^ HFD and n=7 p21^+/TertCi^ HFD mice) C. Left, Insulin Tolerance Test (ITT) was performed by an intraperitoneal injection of insulin (0.3 UI/kg) and glucose was measured via tail clip (Caresens^®^ N, DinnoSanteTM) at different time points. Right, area under the curve (AUC) of the ITT (n=7 p21^+/-^ SD, n=9 p21^+/-^ HFD and n=13 p21^+/TertCi^ HFD mice) D. Hematoxylin and eosin (H&E) staining of visceral adipose tissue (WAT) from p21^+/-^ SD, p21^+/mCherry^ HFD and p21^+/TertCi^ HFD mice. (n=5 p21^+/-^ SD, n=13 p21^+/-^ HFD and n=12 p21^+/TertCi^ HFD mice) E. Adipocyte cell density in the images of H&E-stained WAT was quantified using AdipoSoft ImageJ software (n=5 p21^+/-^ SD, n=13 p21^+/-^ HFD and n=12 p21^+/TertCi^ HFD mice) F. EdU staining quantification into APSC from male WAT from p21^+/-^ SD, p21^+/-^ HFD and p21^+/TertCi^ HFD mice (3 weeks of HFD) after 48 hours pulses of EdU (n=4 p21^+/-^ SD, n=5 p21^+/-^ HFD and n=4 p21^+/TertCi^ HFD mice) G. Expression of the genes involved in lipogenesis, lipolysis and fatty acid oxidation was measured by RT-qPCR in WAT of the p21^+/mCherry^ SD, p21^+/-^ HFD and p21^+/TertCi^ HFD mice (n=5 p21^+/-^ SD, n=6 p21^+/-^ HFD and n=6 p21^+/TertCi^ HFD mice) H. 8-Oxo-désoxyguanosine (oxo-dG) staining of visceral adipose tissue (WAT) from p21^+/-^ SD, p21^+/-^ HFD and p21^+/TertCi^ HFD mice. (n=3 p21+/- SD, n=4 p21+/- HFD and n=4 p21^+/Tert^ HFD mice) I. Quantification of the Oxo-dG positive cell (n=3 p21+/- SD, n=4 p21+/- HFD and n=4 p21^+/Tert^ HFD mice) J. Oxygen consumption rate analyzed by Seahorse in ASPC cell cultures established from WAT of the p21^+/-^ SD, p21^+/-^ HFD and p21^+/TertCi^ HFD mice (n=3 p21^+/-^ SD, n=4 p21^+/-^ HFD and n=4 p21^+/Tert^ HFD mice in duplicate) K. Representative photographs showing adipocytes formation from pre-adipocytes in cell culture L. Expression of the genes involved in lipogenesis was measured by RT-qPCR in pre-adipocytes cell cultures established from WAT of the p21^+/-^ SD, p21^+/-^ HFD and p21^+/TertCi^ HFD mice (n=3 p21^+/-^ SD, n=4 p21^+/-^ HFD and n=5 p21^+/TertCi^ HFD mice) M. SA-β-gal staining of cultured pre-adipocytes from the WAT of the p21^+/-^ SD, p21^+/-^ HFD and p21^+/TertCi^ HFD mice (n=3 p21^+/-^ SD, n=4 p21^+/-^ HFD and n=4 p21^+/TertCi^ HFD mice) N. Quantification of SA-β-gal positive cells in the cultures established from WAT of the indicated groups of mice (n=3 p21^+/-^ SD, n=4 p21^+/-^ HFD and n=4 p21^+/TertCi^ HFD mice) O. mCherry signal was measured by FACS. The percentage of mCherry positive cells is shown in the indicated populations of the SVF. (n=5 p21^+/-^ SD, n=4-7 p21^+/-^ HFD and n=5-12 p21^+/TertCi^ HFD mice) P. p21 mRNA levels was measured by RT-qPCR in stromal vascular fraction (SVF) (n=5 p21^+/-^ SD, n=6 p21^+/-^ HFD and n=6 p21^+/TertCi^ HFD mice) Q. Venn diagram showing the number and overlap of significantly (<0.05) up-regulated genes between p21^+/-^ and p21^+/Tert^ (or p21^+/TertCi^) HFD mice. Gene ontology (GO) enrichment analysis of up-regulated genes in p21^+/Tert^ HFD only, p21^+/TertCi^ HFD only and both, compared to p21^+/-^ HFD mice. R. Venn diagram indicating the number and overlap of significantly (<0.05) down-regulated between p21^+/-^ and p21^+/Tert^ (or p21^+/TertCi^) HFD mice. Gene ontology (GO) enrichment analysis of down-regulated genes in p21^+/Tert^ HFD only, p21^+/TertCi^ HFD only and both, compared to p21^+/-^ HFD mice. All data are shown as the mean ± SEM. *p<0.05 vs. p21^+/-^ SD mice (white bars), #p<0.05 vs. p21^+/-^ HFD (grey bars). Student’s t test or one-way ANOVA with Fisher multiple comparison test.

Moreover, in contrast to our findings in p21^+/Tert^ mice mTERT^Ci^ expression does not attenuate oxidative damage in the AT of HFD-fed p21^+/TertCi^ mice, indicating that telomerase catalytic activity is required to attenuate oxidative damage in the AT of HFD-fed mice (Figure 8H-I). In line with the latter result, we found that preadipocytes isolated from obese p21^+/TertCi^ mice exhibited mitochondrial dysfunction illustrated by decreased oxygen consumption (Figure 8J), suggesting that preservation of mitochondrial function depends on telomere integrity (Sahin *et al*, 2011; Amano *et al*, 2019). Along the same line, we observed that *in vitro* differentiation of ASPCs in mature adipocytes was reduced in p21^+/mCherry^ and p21^+/TertCi^ obese mice compared to pre-adipocytes isolated from SD-fed p21^+/mCherry^ mice (Figure 8K-L). Finally, *in vitro* we observed that preadipocytes derived from p21^+/TertCi^ obese mice exhibited higher levels of senescence than preadipocytes from p21^+/Tert^ obese mice (Figure 8M-N).

Despite these differences in the ability of mTERT and mTERT^Ci^ to generate mature adipocytes, we found that the HFD-induced accumulation of mCherry-positive cells was reduced in p21^+/TertCi^ obese mice as it was in p21^+/Tert^ mice in most of the SVF cell types (Figure 8O) in agreement with the global reduction of p21 expression in the SVF (Figure 8P). Thus, as we previously reported in the lung (Lipskaia *et al*, 2024), mTERT expression in the WAT significantly reduces the fraction of p21-positive cells independently of its catalytic activity. Consistent with this observation, we report that the expression of either mTERT or mTERT^Ci^ attenuates the upregulation of all these genes induced by HFD (Figure 8Q-R). In particular, we found that a number of events related to immune system activation in the adipose tissue of HFD-fed mice are attenuated in p21^+/Tert^ and p21^+/TertCi^ mice.

Thus, expression of mTERT and mTERT^Ci^ have both similar and different effects when expressed under the control of the p21 promoter in the WAT of obese mice.

### mTERT non-canonical function is responsible for remodeling of the macrophage landscape in the AT of obese mice

Our RNA-seq analysis revealed that p21-driven expression of mTERT or mTERT^Ci^ reduced p21 expression and inflammation induced by HFD in the adipose tissue in both p21^+/Tert^ and p21^+/TertCi^ obese mice (Figure 8Q-R). We thus focused our attention on the fate of macrophages and monocytes in the different groups of mice in snRNA-seq data. Macrophages were divided in groups using unsupervised clustering and annotated using known markers (Magalhaes *et al*, 2021; Daemen & Schilling, 2020) (Suppl Figure 8) resulting in 7 classes: monocytes, monocytes derived macrophages, macrophages M0, resident macrophages, intermediate macrophages, proliferating macrophages and macrophages lipid-laden 1 and 2 (Figure 9A). With regard to the proportion of macrophages, p21^+/mCherry^ HFD fed mice showed an increase in intermediate macrophages, Lipid-Laden 1 and 2 macrophages and a decrease in monocytes and resident macrophages suggesting a dynamic shift from monocytes via intermediate to activated lipid laden macrophages. Strikingly, mTERT and mTERT^Ci^ expression under the control of the p21 promoter significantly reduced the number of intermediate macrophages and lipid-laden macrophages, while monocytes, M0 macrophages and resident macrophages were increased (Figure 9B-C). The Lipid-Laden 1 and 2 macrophages represent 42% of the total macrophages in p21^+/mcherry^ obese mice while this proportion dropped to 24% and 19,6% in the AT of p21^+/Tert^ and p21^+/TertCi^ obese mice, respectively (Dataset 2). On the other hand, the proportion of resident macrophages rose from 11% in p21^+/mcherry^ obese mice to 36% and 40% in p21^+/Tert^ and p21^+/TertCi^ obese mice, respectively (Figure 9C and Dataset 2). We inferred from these results that a non-canonical function of mTERT remodels the macrophage landscape in the AT of p21^+/Tert^ and p21^+/TertCi^ obese mice thereby reducing the immune response (Weisberg *et al*, 2003; McNelis & Olefsky, 2014; Chakarov *et al*, 2022; Amano *et al*, 2014).

**Figure 9.**
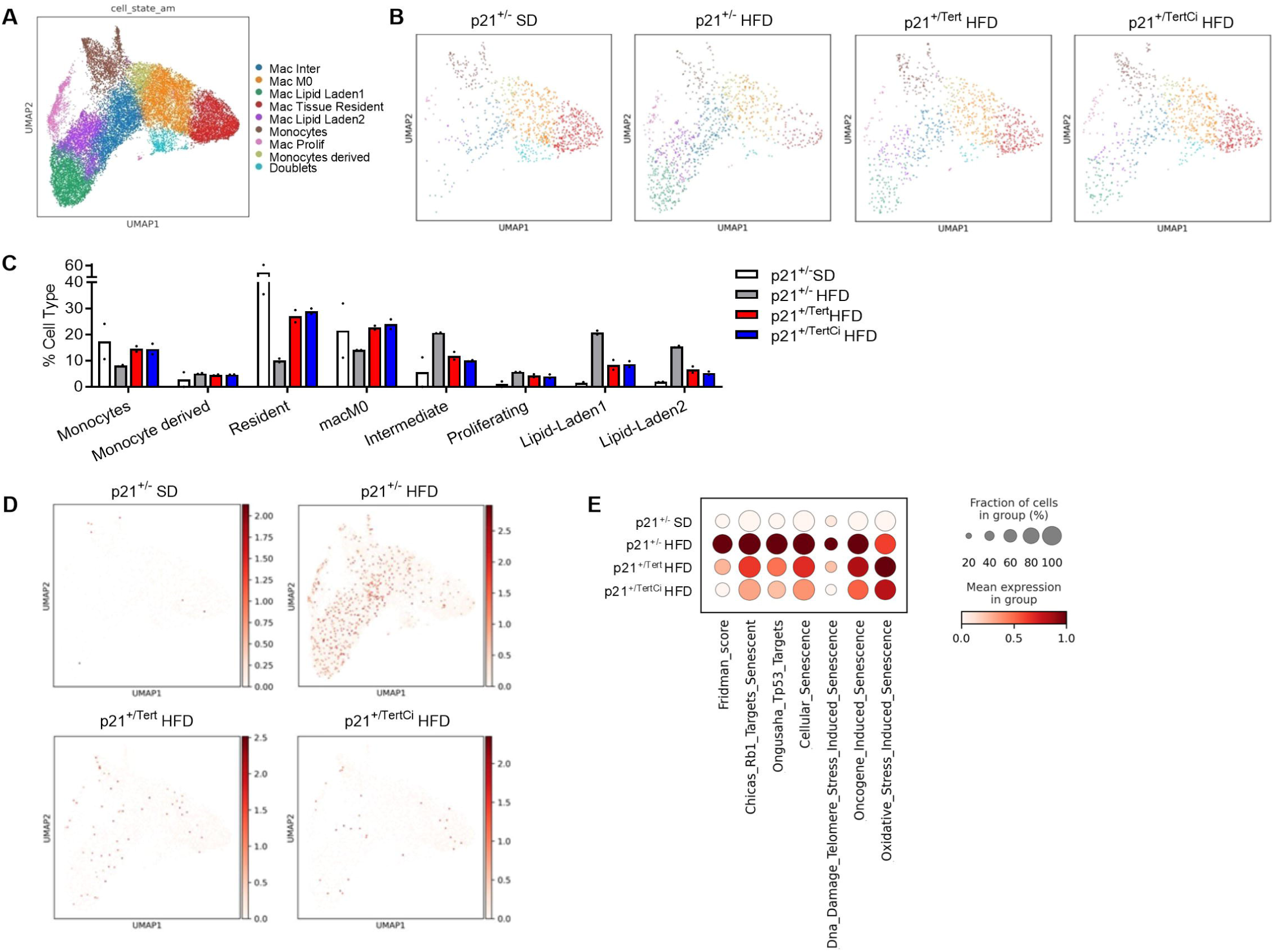
mTERT and mTERTCi remodel the macrophage landscape in the AT of obese mice. A. Subclustering of myeloid cells from snRNAseq data B. Representative myeloid cell subtypes from the WAT of p21^+/-^ SD, p21^+/-^ HFD, p21^+/Tert^ HFD and p21^+/TertCi^ HFD mice C. Mean percentage of myeloid subtypes per sample (excluding doublets) D. p21 expression in different subtypes of myeloid cells in p21^+/mCherry^ SD, p21^+/mCherry^ HFD, p21^+/Tert^ HFD and p21^+/TertCi^ HFD mice E. Scoring of senescence pathways in the different groups of mice in myeloid cells

We used the same approach as for ASPC (we sum the data from the 2 single nucleus RNA-seq experiments, which themselves consist of the average of thousands of values) to determine differences in gene expression within the overall macrophage population in the AT of obese mice. We compared p21^+/Tert^ (and p21^+/TertCi^) mice to p21^+/mCherry^ mice fed with HFD (Dataset 4). As shown in the volcano plots (Suppl Figure 9A-B), many common genes are differentially expressed in macrophages from p21^+/Tert^ and p21^+/TertCi^ obese mice. Many of the down-regulated genes in p21^+/Tert^ and p21^+/TertCi^ obese mice are up-regulated when p21^+/mCherry^ mice are shifted from SD to HFD, whereas up-regulated genes are down-regulated in obese p21^+/mCherry^ mice consistent with the fact that mTERT and mTERT^Ci^ modifies the fate of macrophages in the AT of obese mice.

We then examined p21 expression in different macrophage subpopulations from p21^+/mCherry^ obese mice. We found an increased level of p21 transcripts mainly in monocyte-derived macrophages, intermediate macrophages and lipid-laden macrophages 1 and 2 (Figure 9D). This increase in p21 levels was greatly reduced in p21^+/Tert^ and p21^+/TertCi^ mice (Figure 9D). As for the ASPC, we observed the activation of senescence-related pathways in macrophages from p21^+/mCherry^ HFD mice, an effect that was alleviated in p21^+/Tert^ and counterintuitively even more in p21^+/TertCi^ obese mice (Figure 9E).

Finally, we analyzed in which macrophage subtypes the DEG in macrophages from p21^+/Tert^ and p21^+/TertCi^ obese mice were expressed. We found that genes identified as down-regulated in AT macrophages from p21^+/Tert^ and p21^+/TertCi^ mice were predominantly expressed in Lipid-Laden 1 and 2 macrophages, proliferating, and intermediate macrophages (Ctsl, Igf2r, Znarb3, Pchd7, Fat3, Fgfr2 etc..) (Suppl Figure 9C). Conversely, genes identified as up-regulated (P3h2, Gfra2, Lifr, Aff3, Tsn1, etc..) were preferentially expressed in macrophages M0 and tissue resident macrophages (Suppl Figure 9C). These results confirmed that the ectopic expression of mTERT and mTERT^Ci^ modifies the proportion of macrophage populations in the AT of mice fed with HFD. Of note, the multifaceted transcription factor Zbtb16 (PLZF) (Liu *et al*, 2016) was overexpressed in both ASPCs and macrophages from p21^+/Tert^ and p21^+/TertCi^ HFD mice, with higher expression in p21^+/TertCi^ cells (Dataset 3 and Dataset 4). Zbtb16 retains our attention because it was reported to limit inflammation upon infection in macrophages by stabilizing a co-repressor complex (Sadler *et al*, 2015) and to directly repress transcription of *Cdkn1a*^p21^ by binding to proximal Sp1-binding GC-box 5/6 and distal Tp53-responsive elements of the p21 promoter (Choi *et al*, 2014).

## Discussion

Obesity is regarded as a disease of excess AT leading to metabolic disorders (Hall & Kahan, 2018). However, risk of metabolic dysfunction in obesity appears to be determined by limited capacity for AT remodelling (Sun *et al*, 2011; Shao *et al*, 2018). Telomere shortening-induced senescence has emerged as an important contributor to AT remodeling dysfunction during obesity (Liu *et al*, 2020). Indeed, senescent adipocyte progenitors lose their self-renewal capacity and differentiation potential, contributing to hypertrophic adipocytes formation and metabolic complications in mice and humans (Sakers *et al*, 2022; Gao *et al*, 2020; Conley *et al*, 2020). Cellular senescence, in addition to being a player in AT dysfunction (Sun *et al*, 2011), is above all a hallmark of aging (López-Otín *et al*, 2013; Hernandez-Segura *et al*, 2018), conferring the status of “aging-like” pathology to obesity.

Here, we report that adipose cell progenitors from obese mice display telomere shortening, high p21 expression, increased SA-β-galactosidase activity, decreased adipogenic and differentiation potentials, and mitochondrial dysfunction. We show that the p21 promoter-dependent expression of mTERT in HFD fed mice is associated with reduced p21 expression in particular in ASPCs and macrophages, prevention of telomere attrition, decrease of DNA oxidative damages, and reduced senescence. This facilitates expansion and differentiation of stem and progenitor cells into mature adipocytes as shown by *in vivo* and *in vitro* experiments. Finally, TERT remodels the maladaptive macrophage landscape by preventing polarization toward the inflammatory type that occurs in HFD fed mice. We propose that all these changes, in particular enhanced adipocyte formation, play an important role in improving insulin-resistance and reducing glucose intolerance in p21^+/Tert^ mice fed on a high-fat diet.

The catalytic function of TERT in telomere elongation that ensures long-term cell proliferation has been recognized for many years (Chakravarti *et al*, 2021; Ségal-Bendirdjian & Geli, 2019). However, numerous studies revealed that expression of telomerase has additional roles unrelated to its ability to elongate telomeres (Ségal-Bendirdjian & Geli, 2019) in particular in stem cells (Sarin *et al*, 2005), in progenitor cells (Shkreli *et al*, 2011; Montandon *et al*, 2022), and also in differentiated cells (Lin *et al*, 2018; Neuhöfer *et al*, 2021). TERT was proposed to modulate in the context-dependent fashion critical intracellular signaling pathways such as MYC, WNT, and NF-kB (Park *et al*, 2009; Choi *et al*, 2008; Ghosh *et al*, 2012). We recently reported that mTERT expressed from the p21-promoter promotes the maintenance of high numbers of cells positive for Cd34, a stem cell marker, endowed with proliferative capacities in the lung of old mice (Lipskaia *et al*, 2024). In this context, the question arises as to how mTERT expression promotes adipocyte formation in obese mice fed HFD for 2 months? In obesity, expansion of the AT is driven by increase in number and in size of adipocytes. From a metabolic point of view, adipocyte hyperplasia is more favorable than adipocyte enlargement (Marcelin & Clément, 2021). We show that after 8 weeks on HFD, the AT from obese mice expressing mTERT contains a greater number of adipocytes and of smaller in size compared to control mice. Remarkably, we show that after 3 weeks in HFD, the number of dividing cells within the AT is 2 times greater in p21^+/Tert^ mice than in p21^+/mcherry^ mice. Moreover, *in vivo* and *in vitro* experiments indicate formation of functional adipocytes is promoted by mTERT. In cultured pre-adipocytes from obese mice, p21 promoter-driven expression of mTERT reduces senescence and alleviates mitochondrial dysfunction. Of note, in a previous study (Sun *et al*, 2019), a transient induction of *Tert* expression and telomerase activity was detected during pre-senescence stage of the fibroblasts with shortened telomeres obtained from the progenies of late-generation *Tert*^+/−^ breeding. Transient activity of telomerase observed during pre-senescence of the *Tert*^+/+^ fibroblasts altered the dynamics of DNA Damage Response and delayed senescence relative to the *Tert*^-/-^counterparts (Sun *et al*, 2019). In the same vein, it has very recently been shown that that specific loss of *Tert* in endothelial cell induces senescence in part through a mechanism independent of telomere length that impairs mitochondrial function (Gao *et al*, 2024) supporting a TERT non-canonical function counteracting cell senescence.

We propose that in HFD-fed mice, mTERT by inhibiting p21 expression and senescence pathways facilitates the exit of adipose stem cells from quiescence (de Morree & Rando, 2023) and promote their further expansion. The reduction in oxidative DNA damage in mTERT expressing cells may underlie the reduced activation of signaling pathways leading to p21 expression. Proliferation of ASPCs may then lead to an accumulation of very short telomeres, probably due to replicative stress, that will be repaired by catalytically active mTERT. At a later stage, mTERT maintains a healthy pool of adipocyte precursors which are committed to differentiation into adipocytes. Our results are consistent with a recent study from Wang and colleagues that showed that specific elimination of p21^high^ cells prevents insulin resistance in obese mice. Similar to our observations, they reported that p21^high^ cells in AT were mainly preadipocytes and macrophages (Wang *et al*, 2021).

The second question is how TERT expression promotes the differentiation of ASPCs into mature adipocytes. Changes in the expression of a number of genes in the ASPC of the AT mice could explain the specific features of the p21^+/Tert^ ASPC. For instance, we found that Fgfr2 that is known to be a driver of osteoblast differentiation and expressed in adipocytes (Hutley *et al*, 2011; Du *et al*, 2012) is down regulated in p21^+/Tert^ ASPCs. Other genes playing a role in bone formation (Postn, Gla, Aebp1, Adam12, Runx2) as well as a number of cathepsin and related proteins (Ctsl, Ctsb, Ctsd, Ctsp, Ctss) that promote osteocyte differentiation (Choi *et al*, 2023) were found also to be down regulated (Dataset 1) (Suppl Figure **10** and Suppl Figure **11**). These changes are anticipated to promote ASPCs commitment to adipogenic differentiation. Along the same line, we found that the conditional expression of mTERT promotes the up-regulation of a number of genes (Cav1 / Dact1 / Foxo1 / Gli3 / Gpc3 / Mllt3 / Ppp2r3a / Ror2 / Shisa6 / Sox13 / Tmem64 / Wnt5b) (Dataset 3) described as negative regulators of the Wnt /β-catenin signaling pathway. Because Wnt signaling preserves progenitor cell multipotency during the early stage of adipogenesis while its subsequent inactivation is necessary for ASPSc differentiation into adipocytes (Park *et al*, 2009; Palani *et al*, 2023), it is conceivable that mTERT promotes ASPCs commitment to the adipocyte differentiation by reducing Wnt signaling. Another important piece of information lies in the fact that expression of p21 and senescence-associated genes is reduced in ASPCs from the AT of p21^+/Tert^ HFD mice. Lower p21 level in p21^+/Tert^ HFD obese mice is expected to promote the activity of Cdk2 and Cdk4 that has been reported to favor clonal expansion and differentiation of adipocyte precursors (Abella *et al*, 2005).

How mTERT expression regulates (directly or indirectly) gene expression in ASPC? Very recently, it has been reported that conditional deletion of mTERT in spermatogonial stem cells impedes the formation of competitive clones (Hasegawa *et al*, 2024). It is proposed that mTERT favors chromatin accessibility across many genes which expression promotes competitive clone formation. Interestingly, the reduction in open chromatin in mTERT-KO spermatogonial stem was particularly evident in Zbtb16 (Hasegawa *et al*, 2024) that is overexpressed in ASPCs from p21^+/TertCi^ HFD mice. Along the same line, human colorectal tumor samples expressing high levels of TERT show increased levels of open chromatin at iron metabolism related genes compared to tumor samples expressing low levels of TERT (Shanmugam *et al*, 2024). These studies suggest that mTERT may act directly at the chromatin level. Interestingly, the Abca8b, Abca8a, Abca9 and Abca6 genes of the ABC (ATP-binding-cassette) transporter subfamily, which form a cluster on chromosome 1, are upregulated in p21^+/Tert^ ASPCs (and other cell types). These genes are all transcribed in the same direction and two strong CTCF binding sites bracket them suggesting that these genes could form a single chromatin loop extruded by CTCF/cohesin. Along the same line, *ABCC5AS1 and THSD7A*, among the top upregulated genes in p21^+/Tert^ ASPCs, exhibit a strong correlation between their methylation state and expression in mesenchymal cells from visceral adipose tissue of obese patients (Mirzaeicheshmeh *et al*, 2021). It is tempting to speculate that mTERT could interact with chromatin remodelers and/or epigenetic modifiers to regulate genes, even over long distances. Alternatively, we may also envision that mTERT may directly or indirectly regulate phosphatase and/or kinase activities. This assumption is supported by very recent results showing that activation and clonal expansion of kidney podocyte progenitors induced by the Dox-induced expression of mTERT^Ci^ is suppressed by the inactivation of Wip1 (Ppm1d) (Duret *et al*, 2024), a Ser/Thr phosphatase of the PP2C family that is crucial for the deactivation of several signalling pathways (Nahta & Castellino, 2021) including the DNA Damage Response (Shreeram *et al*, 2006).

In this study, we wondered which effects associated with TERT conditional expression were independent of its catalytic activity. To this end, we expressed a catalytically inactive TERT under the control of the p21 promoter. Both *in vivo* and *in vitro*, our results indicate that the formation of functional adipocytes is mainly enhanced by mTERT, but not by mTERT^Ci^, indicating that the catalytic activity of mTERT is important for the generation of mature adipocytes. We also found that telomerase catalytic activity in p21^+/Tert^ mice was required to improve mitochondrial function in preadipocytes cultured in vitro and for the reduction of oxidative damage in the AT of obese mice. The improvement in mitochondrial activity does not appear to be linked to variations in the expression of the seven sirtuins previously described as reduced in the liver of mice with dysfunctional telomeres (Amano *et al*, 2019). Indeed, expression of sirtuins does not appear to vary in the ASPCs of the AT from obese mice of the 3 phenotypes (p21^+/mCherry^, p21^+/Tert^ and p21^+/TertCi^). The maintenance of functional telomeres therefore appears to be necessary for steps required for the formation of mature adipocytes, presumably for the clonal expansion of ASPCs as discussed above. These results are consistent with the fact that TERT has a better ability than TERT^Ci^ to reduce metabolic syndromes in obese mice.

However, in agreement with our previous study (Lipskaia *et al*, 2024), we found that both p21^+/Tert^ and p21^+/TertCi^ mice have reduced p21 expression in the WAT under the stress of overfeeding. The results also indicate that mTERT and mTERT^Ci^ conditional expression show similar transcriptomic signatures in ASPCs of obese mice. Consequently, TERT’s canonical and non-canonical functions contribute to the effects mediated by TERT. TERT’s non-canonical functions are likely to be required for the attenuation of the metabolic defects induced by HFD, but not sufficient.

Strikingly, our study also reveals that a function of TERT independent of its catalytic activity modifies the macrophage distribution in the AT of p21^+/Tert^ and p21^+/TertCi^ mice fed with HFD. We observe in the AT from p21^+/mTert^ and p21^+/TertCi^ obese mice a reduction in the number of intermediate macrophages and lipid-laden macrophages, while monocytes, M0 macrophages and resident macrophages are increased. This is in particular reflected by the reduced Trem2 signature (Trem2, Ctsl, Ctsb, Lgals1, Lagals3, Spp1, ApoE,…) that characterizes lipid associated macrophages (Sárvári *et al*, 2021). Because Trem2 is a major macrophage sensor of extracellular lipids (Sárvári *et al*, 2021), it is possible that the altered macrophage landscape is a consequence of the reduced amount of pathological lipids in the AT from p21^+/Tert^ and p21^+/TertCi^ obese mice. It remains to be determined whether the change in the macrophage landscape is a consequence of the intrinsic action of mTERT in macrophages, or results from the reduced oxidative damages and senescence in preadipocytes, or a synergistic effect of both.

It has recently been shown that p21 expressed in cells under oncogenic/genotoxic stress activates a retinoblastoma protein (Rb)-dependent transcriptional program involving specific SMAD and STAT transcription factors, leading to the expression of a number of secreted factors (called PASP) including CXCL14 that rapidly recruits macrophages. If activation is sustained, secreted factors polarize macrophages towards an inflammatory phenotype (Sturmlechner *et al*, 2021). The single nucleus RNA-seq data of this study shows that senescence activation programs mediated by Rb or Tp53 are alleviated in many cell types of the p21^+/Tert^ and p21^+/TertCi^ AT. Although, we do not find that Cxcl14 is differentially expressed in the SVF fraction of the AT from the different groups of obese mice, expression of a number of chemokines attracting macrophages such as C3ar1, C5ar1, and Ccl2 is reduced in the p21^+/Tert^ and p21^+/TertCi^ SVF. This could account for the changes in AT macrophage landscape. We found that many of the most down-regulated genes in p21^+/Tert^ and p21^+/TertCi^ macrophages were predominantly expressed in mac_lipid_laden 1 and 2 and in proliferating macrophages. Interestingly, many of these down-regulated genes (Fat3, Igf2r, Pcdh7, Fth1, Ftl1, Fgfr2, Ctsl, Trem2, ApoE) were also down-regulated in ASPC suggesting common mechanisms of gene regulation induced by mTERT expression in both cell types. Interestingly, conditional macrophage-specific Fth1 knockout was reported to alleviate obesity and diabetes in mice fed with HFD (Ikeda *et al*, 2020). In contrast, upregulated genes were predominantly expressed in tissue-resident macrophages and somehow reflect the predominance of these type of macrophages in the adipose tissue of p21^+/Tert^ and p21^+/TertCi^ mice.

Importantly, our results indicate that it is mainly males that are metabolically affected by HFD in the short-term (here 8 weeks), while females are protected against HFD-induced insulin resistance and glucose intolerance (Pettersson *et al*, 2012). Many studies have attributed this protection to estrogen, since older postmenopausal female mice are no longer protected against obesity (Riant *et al*, 2009; Zhu *et al*, 2013). In our study, obese females did not show higher expression of p21 in adipose tissue than lean females, suggesting that female mice are better protected in particular against HFD-induced p21-expression in the AT. Whether this is due to a better protection to DNA damage remains to be determined. Gao and colleagues also found that clearance of p21^high^ cells had less metabolic benefits in female than male obese mice after 2 months HFD (Gao *et al*, 2020). These results support the idea that female, somehow, are better protected against senescence and may therefore explain sex-differences in senescence associated diseases.

Taken together, these results support the idea that the conditional expression of TERT in pre-senescent cells (p21^high^ cells) could be used as a therapeutic tool to regulate adipose tissue remodeling and insulin-resistance in obesity. TERT gene therapy in aged mice has been shown to delay aging without increasing the number of cancers (Bernardes de Jesus *et al*, 2012). Very recently, it was reported that administration of a transcriptional activator of TERT in aged mice alleviates cellular senescence and systemic inflammation in different organs opening new avenues for therapeutic interventions (Shim *et al*, 2024).

## Materials and Methods

### Animals

Mice p21^+/mCherry^, p21^+/mTert^ and p21^+/mTertCi^ were housed under controlled conditions of temperature (21±1°C), hygrometry (60±10%) and lighting (12 h per day). Animals were acclimatized in the laboratory for one week before the start of the experiments. Mice were fed either a standard diet SD (A04, SAFE Diet, Augy, France) or a high fat diet HFD (60 kcal % fat, SAFE Diet, Augy, France). All animals received care according to institutional guidelines, and all experiments were approved by the Institutional Ethics committee number 16, Paris, France (licence number 16-090). During follow-up, animals underwent body-weight, metabolic assessments. For EdU (EdU kit reagents Invitrogen C10646) treatments, mice were given two intraperitoneal injections of 5mg/kg EdU in sterile PBS every 12 hours two days before the sacrifice. Mice were euthanatized and organs and blood were collected and processed for further evaluations.

### Fasting blood glucose, glucose and insulin tolerance tests

Whole-body glucose tolerance and insulin sensitivity were assessed in all groups at weeks 12^th^ and 13^th^ by intraperitoneal glucose (GTT) and insulin (ITT) tolerance tests, respectively. First, blood was obtained via tail clip to assess fasting blood glucose (Caresens^®^ N, DinnoSanteTM). Then, mice received either glucose (1.5 g/kg) or insulin (0.3 UI/kg) solution (sigma I9278) by intraperitoneal injection, and blood glucose was measured at 15, 30, 60, 90 and 120 min after the injection. The area under the curve (AUC) for the glucose excursion was calculated using Graphpad Prism.

### Plasma

Enzyme-linked immunosorbent assay (ELISA) kit (Life Technology Ref KMC2281) was used to measure Leptin level in the blood plasma.

### Analysis of mRNA expression

For RNA extraction the samples were lysed with Qiazol (Qiagen, France) in the presence of chloroform, and total RNA was purified on mini-columns using RNeasy extraction kit (Qiagen, ref 74104). First-strand cDNA was synthesized from total RNA using the High-Capacity cDNA Reverse Transcription Kit (Life Technologies, ref 4368814). Quantitative real-time PCR (qPCR) was performed in a StepOnePlus Real-Time PCR. Gene expression was assessed by the comparative CT (ΔΔCT) method with β-actin as the reference gene.

### RNA Sequencing and Data Analysis

Quality control of samples. RNA sequencing was performed by Novogene company (Cambridge, UK). RNA integrity was assessed using the RNA Nano 6000 Assay Kit of the Bioanalyzer 2100 system (Agilent Technologies, CA, USA). Library preparation for Transcriptome sequencing. Total RNA was used as input material for the library preparations. Briefly, mRNA was purified from total RNA using poly-T oligo-attached magnetic beads. Fragmentation was carried out using divalent cations under elevated temperature in First Strand Synthesis Reaction Buffer (5X). First strand cDNA was synthesized using random hexamer primer and M-MuLV Reverse Transcriptase (RNase H-). Second strand cDNA synthesis was subsequently performed using DNA Polymerase I and RNase H. Remaining overhangs were converted into blunt ends via exonuclease/polymerase treatment. After adenylation of the 3’ ends of DNA fragments, Adaptor with hairpin loop structure was ligated to prepare for hybridization. In order to select cDNA fragments in the reange of 370-420 bp, the library fragments were purified with AMPure XP system (Beckman Coulter, Beverly, USA). Then PCR was performed with Phusion High-Fidelity DNA polymerase, using the Universal and Index (X) primers. At last, PCR products were purified (AMPure XP system) and library quality was assessed on the Agilent Bioanalyzer 2100 system. Clustering and sequencing. The clustering of the index-coded samples was performed on a cBot Cluster Generation System using TruSeq PE Cluster Kit v3-cBot-HS (Illumia) according to the manufacturer’s instructions. After cluster generation, the library preparations were sequenced on the Illumina Novaseq platform generating 150 bp paired-end reads.

Sequencing quality control was determined using the FastQC tool (http://www.bioinformatics.babraham.ac.uk/projects/fastqc/) and aggregated across samples using MultiQC (v1.7) (Ewels *et al*, 2016). Reads with a Phred quality score of less than 30 were filtered out. Reads were mapped to a customized mouse mm10 genome (containing the mCherry transgene) using Subread-align (v1.6.4) (Liao *et al*, 2013) with default parameters. Gene expression was determined by counting mapped tags at gene levels using featureCounts (v1.6.4) (Liao *et al*, 2014), and differentially expressed genes (DEG) were identified using DESeq2 (v1.26.0) (Love *et al*, 2014). Statistical significance was inferred at P<0.05 (Benjamini-Hochberg corrected).

Gene Ontology (GO) enrichment analysis was performed on DEGs using clusterProfiler (v4.6.0) (Wu *et al*, 2021) and considering a background. Only the biological process (BP) category was chosen and GO terms with corrected P value less than 0.05 were considered significantly enriched. Enrichment heatmaps were generated using heatmap.2 (v<u>3.1.3.1)</u> (Warnes *et al*, 2009), highlighting the 10 most differentially regulated genes per GO term.

String analysis (https://string-db.org) was performed to investigate interactions between differentially expressed gene.

### Single nucleus RNA sequencing

Single nucleus sequencing was carried out using visceral WAT (snap frozen and stored at -80C) in n=2 mice per group. WAT samples were processed in batches of n=2, randomizing the order, to minimize between condition batch effects. Nuclei extraction and isolation was done using a modified version of a previously described protocol (Nee *et al*, 2018). Briefly, frozen WAT (approx. 200mg) was cut into small pieces <0.2cm and homogenized with ice cold lysis buffer (Tris-HCL 10mM, NaCl 10mM, MgCl2 3mM, 0.1% NP40, 0.2U/µl RNase inhibitor) in a glass dounce homogenizer. Following homogenization, the sample was filtered through a 100µM cell strainer and centrifuged at 500 x g and 4oC for 5mins. After lipid layer and supernatant removal, the nuclei pellet was resuspended in wash buffer (PBS + 1% BSA and 0.2U/µl RNase inhibitor) and recentrifuged at 500 x g and 4°C for 5mins. Isolated nuclei were then resuspended in wash buffer with DAPI (0.1µg/ml) and filtered through 35µM cell strainer into a FACS tube. Stained nuclei were sorted by flow cytometry using the BD FACS Aria III (85-microm nozel) to remove excess lipid/debris and clumped nuclei, applying the following sequential gates: i. FSC-A and SSC-A; ii. FSC-H and FSC-A; iii. FSC-A and 450/40V-A. Nuclei were sorted into cold (4C) BSA and RNase-inhibitor rich collection buffer to prevent re-clumping and RNA degradation; final nuclei suspension post-FACS concentrations (BSA ∼2% and RNase inhibitor ∼0.3U/µl).

An estimated ∼5-10K nuclei per sample were loaded onto Chromium Next GEM Single Cell 3’ Chips for GEM generation and barcoding, followed by clean up, cDNA amplification (10-12 cycles) and library construction, according to 10X manufacturer instructions. Single Cell 3ʹ Gene Expression libraries were quantified using the Agilent Bioanalyzer (High Sensitivity chip), and pooled for sequencing (Nextseq2000), aiming for a final sequencing depth of >20K reads/called-cell/sample. Sequenced libraries were then de-multiplexed and analysed using CellRanger v5.0.1 and bcl2fastq v2.20.0. Individual libraries were de-multiplexed with CellRanger mkfastq (allowing 1 mismatches) and the CellRanger count pipeline was used to align to the mouse genome (customized with mouse genotype alleles) and count UMIs. UMIs QC (i.e. ambient noise versus cell signals filtering) was carried out using barcodeRanks function within DropletUtils package (v1.18.0) (Lun *et al*, 2019). Sample level quality control analyses were then carried out using perCellQCMetrics, perFeatureQCMetrics within scater package (v1.26.0) (McCarthy *et al*, 2017). Only the cells exhibiting median-absolute-deviation (MAD) values less than five for the different QC metrics were retained for further analysis. Cell doublets were identified using DoubletFinder package (v2) (McGinnis *et al*, 2019). After quality filtering individual samples were normalized (NormalizeData) and transformed (normalization.method = “LogNormalize”), and then integrated through canonical correlation analysis (CCA) involving FindIntegrationAnchors and IntegrateData functions within Seurat (v4.3.0) (Hao *et al*, 2021) to create a single atlas of n=42343 cells. The integrated dataset was clustered using the Louvain algorithm (resolution = 0.5) through the FindNeighbors and FindClusters functions implemented in Seurat. Cell type inference step was carried out using 3 tools (TransferData within Seurat, ACTINN (Ma & Pellegrini, 2020), and CaSTLe (Lieberman *et al*, 2018)) and an already published single cell references (Emont *et al*, 2022). The final cell type annotation was determined by a democratic approach, where the most frequently assigned cell type across the three rounds of annotation was assigned to each cell.

Each major WAT cell type was then reintegrated in Scanpy using Harmony (Korsunsky *et al*, 2019) and BBKNN (Polański *et al*, 2020), subclustered using the Leiden algorithm implemented within Scanpy, and manually annotated with established marker genes and published single cell references (Magalhaes *et al*, 2021; Daemen & Schilling, 2020) (Details below and in main text comment). To assist annotation, unbiased markers were obtained with the rank_gene_groups function within Scanpy, using “t-test_overestim_var” method with a Benjamini-Hochberg correction. Droplets containing markers of different cell types were further considered doublets and removed from downstream analysis. Given the limited number of samples, gene expression between groups was tested using the nonparametric Wilcoxon rank sum test treating each cell as a unique data point. Statistical significance was inferred at P<0.05 (Bonferroni corrected). Senescence scores were obtained with the “score_genes” function within Scanpy, using published lists and a control size of 50 genes (Subramanian *et al*, 2005).

### Histology and immunohistochemistry

Fresh visceral adipose tissue (WAT) was fixed in 10% phosphate-buffered formalin overnight. Paraffin wax sections of 5 µm were prepared for immunostaining. Images of the haematoxylin-eosin-stained tissues were analyzed using Adiposoft Image J plugin (https://imagej.net/plugins/adiposoft). Click-EdU staining was performed on FFPE slides as described in the Invitrogen kit protocol (Click-EdU kit reagents Invitrogen C10646).

### Stromal vascular fraction (SVF) isolation and culture

Visceral fat was excised from the mice, minced with scissors, and digested for one hour at 37°C in 1 mg/ml collagenase type II digestion buffer (Life Technology, France; 17100017) in sterile Hank’s Balanced Salt Solution (Life Technologies, ref 14185-052) containing 3% bovine serum albumin. After digestion, SVF were separated from adipocytes by centrifugation (300g, 3min) and two filtration steps (70µm and 40µm cell strainers). SVF were plated in a 6-well plate in DMEM supplemented with 0.5% Pen/Strep, 10% new born calf serum and 1% pyruvate.

### Imaging flow cytometry analysis of the SVF

Purified SVF cells were incubated with a panel of antibodies (CD45 BD Bioscience 561047, CD31 BD Bioscience 612802, CD34 BioLegend 119314, CD29 BioLegend-102218, ScaI BioLegend 108127, CD24 BioLegend-101823, PDGFRα BioLegend-135905, F4/80 BioLegend-123115, CD11b BD Bioscience-565976, CD11c BD Bioscience-749038, Live Dead Nir Cytek R7-60008) for 30 min on ice. Cells were then washed with 0.5% FBS/PBS FACS buffer and resuspended in 300 µL of FACS buffer. Cells were detected and their fluorescence was measured using Aurora flow cytometer (Cytek, Amsterdam, Netherland). Data analysis was performed in the SpectroFlow software.

### Telomere Shortest Length Assay (TeSLA)

TeSLA was performed as described previously (Lipskaia *et al*, 2024).

### Cellular Bioenergetic Analysis Using the Seahorse Bioscience XF Analyzer

Bioenergetic profiles of the adipocytes were determined using a Seahorse Bioscience XF24 Analyzer (Billerica, MA, USA) that provides real-time measurements of oxygen consumption rate (OCR), indicative of mitochondrial respiration, and extracellular acidification rate (ECAR), an index of glycolysis as previously described (Braud *et al*, 2018).

### Statistical analysis

Data are expressed as mean values ± standard error of the mean (SEM). Statistical significance was tested using either one or two-way analysis of variance (ANOVA) with Fisher multiple comparison test. The results were considered significant if the p-value was <0.05.

### Data availability

Single-nucleus RNA-seq data and Bulk RNA-seq data have been deposited in the Gene Expression Omnibus (GSE261438 and GSE261439).

## Acknowledgements

We thank Florent Ginhoux and Svetoslav Charakov for helpful advices on macrophage subdivision and biology. We thank Jean-Charles Graziano from CRCM’s animal facility and the CRCM’s Integrative BioInformatics platform (Cibi) and Cytometry platform. We would like to thank Frédéric Fiore (CIPHE, Marseille) for his advice on generating mouse models. Work in VG’s Lab is supported by “La Ligue Nationale Contre le Cancer”, Equipe Labellisée, the “Institut National du Cancer” (INCA, PLBIO 2019), the Agence Nationale de la Recherche (ANR) (Grant THALATEL), the cross-cutting Inserm program on aging (AGEMED) and the Inserm INTERAGING Program. This study was partly supported by research funding from Gefluc and Canceropole Sud. Core support from MRC (MC_U120085810) and CRUK (C15075/A28647) funded this research in J.G’s laboratory. W.S. was funded by the MRC (MC_UP1605/7) and Wellcome Trust (219602/Z/19/Z).

## Author Contributions

L.B. and V.G. designed the study. L.B coordinated most the experiments. L.B, M.B., M.R performed metabolic tests, qPCR, FACS experiments. M.B. and D.C. performed TeSLA experiments. W.S, L.M.A and A.M performed and analyzed the single nucleus RNA sequencing experiment. A.D performed the 8-Oxo-dG experiments. J.G. contributed to design the single nucleus RNA sequencing experiment. VG, W.S, and J.G secured funding. J.V analyzed the RNA sequencing and the single cell sequencing data. L.B and V.G. wrote the manuscript with inputs of W.S. and J.G. L.B and V.G are the guarantors of this work and, as such, have full access to all the data in this study and take responsibility for the integrity of the data and the accuracy of the data analysis.

## Conflict of interests

J.G. has acted as a consultant for Unity Biotechnology, Geras Bio, Myricx Pharma Ltd., and Merck KGaA; owns equity in Geras Bio and share options in Myricx Pharma Ltd. and is a named inventor in MRC and imperial College patents related to senolytic therapies (unrelated to the work described here). J.G.’s lab received Pfizer and Unity Biotechnology unrelated to the work described here.

## Supplemental Figure Legends

**Suppl Figure 1. mTERT under the control of the p21 promoter does not impact female metabolism**

A. Body weight (g) of female mice (n=8 p21^+/-^ SD, n=5 p21^+/-^ HFD and n=8 p21^+/Tert^ HFD mice)
B. Weight of the white adipose tissue (WAT) (g) (n=8 p21^+/-^ SD, n=5 p21^+/-^ HFD and n=7 p21^+/Tert^ HFD mice)
C. Leptin plasmatic level was measured by Leptin Elisa assay (n=6 per group)
D. Glucose Tolerance Test (GTT) and area under the curve of the GTT (n=8 p21^+/-^ SD, n=7 p21^+/-^ HFD and n=6 p21^+/Tert^ HFD mice)
E. Insulin Tolerance Test (ITT) and area under the curve of the ITT (n=4 p21^+/-^ SD, n=7 p21^+/-^ HFD and n=7 p21^+/Tert^ HFD mice)
F. p21 mRNA levels was measured by RT-PCR in stromal vascular fraction (SVF) of male and female (n=8 p21^+/-^ SD male, n=9 p21^+/-^ HFD male and n=7 p21^+/Tert^ HFD male and n=6 p21^+/-^ SD female, n=6 p21^+/-^ HFD female and n=6 p21^+/Tert^ HFD female mice)

Values represent the mean ± SEM. *p<0.05 vs. p21^+/-^ SD mice (white bars), ^#^p<0.05 vs. p21^+/-^ HFD (grey bars). Student’s t test or one-way ANOVA with Fisher multiple comparison test.

**Suppl Figure 2. Adipose tissue stromal vascular fraction (SVF) subpopulation of cells analyzed by FACS**

Gating strategy, Flow cytometry gating strategy used to define Immune Cells (CD45^+^ F4/80^-^), Leukocytes (CD45^+^ F4/80^-^ CD11b^-^ CD11c^-^), Myeloid cells (CD45^+^ F4/80^-^ CD11b^+^ CD11c^-^), Dendritic Cells type 1 (DC) (CD45^+^ F4/80^-^ CD11b^+^ CD11c^+^) and Dendritic Cells type 2 (DC-2) (CD45^+^ F4/80^-^ CD11b^-^ CD11c^+^), Macrophages (CD45^+^, F4/80^+^), MacCD11b^+^ (CD45^+^, F4/80^+^ CD11b^+^ CD11c^-^), MacCD11c^+^ (CD45^+^ F4/80^+^ CD11b^+^ CD11c^+^), Endothelial cells (CD45^-^ CD31^+^), Pericytes (CD45^-^ CD31- CD34^+^ ScaI^-^), Pre-adipocytes (CD45^-^ CD31^-^ CD34^+^ ScaI^+^ CD29^+^ PDGFRα^+^ CD24^-^) and Adipose Stem and Progenitors Cells (ASPC) (CD45^-^ CD31^-^ CD34^+^ ScaI^+^ CD29^+^ PDGFRα^+^ CD24^+^). Dead cells were excluded with Live Dead Nir, cells were gated based on SSC-A versus FSC-A and singlets were selected from the SSC-A versus SSC-H dot plot.

**Suppl Figure 3. Analysis of short telomeres in the WAT**

A. Telomere Shortest Length Assay (TeSLA) of WAT using 30 pg of gDNA for each TeSLA PCR (n=3 per group).
B. Cumulative percentage of Tesla bands length bin (kb) (n=3 per group).
C. Mean telomere length of the telomeres detected by the Tesla method in the indicated groups of mice (n=3 per group).

Values represent the mean ± SEM. *p<0.05 vs. p21^+/-^ SD mice (white bars), ^#^p<0.05 vs. p21^+/-^ HFD (grey bars). Student’s t test or one-way ANOVA with Fisher multiple comparison test.

**Suppl Figure 4. Volcano plots of differentially regulated genes (Related to Figure 3)**

Volcano plots depicting significant (<0.05) differentially expressed genes between p21^+/-^ SD and HFD mice and associated Gene Ontology (GO) analysis revealing enrichment of genes involved in the indicated pathways.

A. Volcano plots depicting significant (<0.05) differentially expressed genes between p21^+/-^ SD and p21^+/-^ HFD mice and associated Gene Ontology (GO) analysis revealing enrichment of genes involved in the indicated pathways.
B. Volcano plots depicting significant (<0.05) differentially expressed genes between p21^+/-^ HFD and p21^+/Tert^ HFD mice and associated Gene Ontology (GO) analysis revealing enrichment of genes involved in the indicated pathways.

**Suppl Figure 5. Markers used for the subclustering of ASPC**

A. Top unbiased markers of the different ASPC subtypes. Dotplots represent scaled expression per gene.
B. Handpicked markers to assist the identification of ASPC subtypes. Dotplots represent scaled expression per gene.

**Suppl Figure 6. p21 expression in AT cell types and senescence signature p21 positive cells**

A. Ratio of p21 positive to p21 negative cells in the different groups of mice
B. Violin plots showing p21 expression in the indicated cell types
C. Score of senescence signatures in all cell types
D. Score of senescence signatures in adipocytes
E. Score of senescence signatures in myeloid cells

**Suppl Figure 7. Volcano plots of DEG in ASPC subpopulation.**

Volcano plots depicting significant (<0.05) differentially expressed genes between p21^+/mCherry^ HFD and p21^+/Tert^ HFD ASPC

**Suppl Figure 8. Markers used for the subclustering of macrophages data**

A. Top unbiased markers of the different macrophage subtypes. Dotplots represent scaled expression per gene.
B. Handpicked markers to assist the identification of macrophage subtypes. Dotplots represent scaled expression per gene.

**Suppl Figure 9. Volcano plots of DEG in macrophages subpopulation**

A. Volcano plots depicting significant (<0.05) differentially expressed genes between p21^+/mCherry^ HFD and p21^+/Tert^ HFD macrophages
B. Volcano plots depicting significant (<0.05) differentially expressed genes between p21^+/mCherry^ HFD and p21^+/TertCi^ HFD macrophages
C. Dotplots show scaled (per gene) mean expression of selected DEG per myeloid subtypes between p21^+/mcherry^ HFD vs p21^+/Tert^ HFD and Dotplots show scaled (per gene) mean expression of selected DEG per myeloid subtypes between p21^+/mCherry^ HFD vs p21^+/TertCi^ HFD

**Suppl Figure 10. Predicted gene interactions within down-regulated genes in p21^+/Tert^ ASPC.**

The interaction network was downloaded from the STRING database. K-Means clustering was used to cluster the genes.

**Suppl Figure 11. Predicted gene interactions within down-regulated genes in p21^+/TertCi^ ASPC.**

The interaction network was downloaded from the STRING database. K-Means clustering was used to cluster the genes.

## Dataset Legends

**Dataset 1. RNA-seq in total SVF and GO**

**Dataset 2. Number of cells for each cell type and for each condition**

**Dataset 3. Gene expression in ASPC**

**Dataset 4. Gene expression in Macrophages**

## References

Abella A, Dubus P, Malumbres M, Rane SG, Kiyokawa H, Sicard A, Vignon F, Langin D, Barbacid M & Fajas L (2005) Cdk4 promotes adipogenesis through PPARγ activation. Cell Metab 2: 239–249

Amano H, Chaudhury A, Rodriguez-Aguayo C, Lu L, Akhanov V, Catic A, Popov YV, Verdin E, Johnson H, Stossi F, et al (2019) Telomere Dysfunction Induces Sirtuin Repression that Drives Telomere-Dependent Disease. Cell Metab 29: 1274–1290.e9

Amano SU, Cohen JL, Vangala P, Tencerova M, Nicoloro SM, Yawe JC, Shen Y, Czech MP & Aouadi M (2014) Local proliferation of macrophages contributes to obesity-associated adipose tissue inflammation. Cell Metab 19: 162–171

Bernardes de Jesus B, Vera E, Schneeberger K, Tejera AM, Ayuso E, Bosch F & Blasco MA (2012) Telomerase gene therapy in adult and old mice delays aging and increases longevity without increasing cancer. EMBO Mol Med 4: 691–704

Braud L, Pini M, Muchova L, Manin S, Kitagishi H, Sawaki D, Czibik G, Ternacle J, Derumeaux G, Foresti R, et al (2018) Carbon monoxide–induced metabolic switch in adipocytes improves insulin resistance in obese mice. JCI Insight 3

Cannavino J & Gupta RK (2023) Mesenchymal stromal cells as conductors of adipose tissue remodeling. Genes Dev 37: 781–800

Chakarov S, Blériot C & Ginhoux F (2022) Role of adipose tissue macrophages in obesity-related disorders. J Exp Med 219: e20211948

Chakravarti D, LaBella KA & DePinho RA (2021) Telomeres: history, health, and hallmarks of aging. Cell 184: 306–322

Chen S, Yeh F, Lin J, Matsuguchi T, Blackburn E, Lee ET, Howard BV & Zhao J (2014) Short leukocyte telomere length is associated with obesity in American Indians: the Strong Heart Family study. Aging 6: 380–389

Choi J, Southworth LK, Sarin KY, Venteicher AS, Ma W, Chang W, Cheung P, Jun S, Artandi MK, Shah N, et al (2008) TERT promotes epithelial proliferation through transcriptional control of a Myc- and Wnt-related developmental program. PLoS Genet 4: e10

Choi J-W, Lim S, Jung SE, Jeong S, Moon H, Song B-W, Kim I-K, Lee S, Hwang K-C & Kim SW (2023) Enhanced Osteocyte Differentiation: Cathepsin D and L Secretion by Human Adipose-Derived Mesenchymal Stem Cells. Cells 12: 2852

Choi W-I, Kim M-Y, Jeon B-N, Koh D-I, Yun C-O, Li Y, Lee C-E, Oh J, Kim K & Hur M-W (2014) Role of promyelocytic leukemia zinc finger (PLZF) in cell proliferation and cyclin-dependent kinase inhibitor 1A (p21WAF/CDKN1A) gene repression. J Biol Chem 289: 18625–18640

Conley SM, Hickson LJ, Kellogg TA, McKenzie T, Heimbach JK, Taner T, Tang H, Jordan KL, Saadiq IM, Woollard JR, et al (2020) Human Obesity Induces Dysfunction and Early Senescence in Adipose Tissue-Derived Mesenchymal Stromal/Stem Cells. Front Cell Dev Biol 8

Daemen S & Schilling JD (2020) The Interplay Between Tissue Niche and Macrophage Cellular Metabolism in Obesity. Front Immunol 10

Du X, Xie Y, Xian CJ & Chen L (2012) Role of FGFs/FGFRs in skeletal development and bone regeneration. J Cell Physiol 227: 3731–3743

Duret LC, Hamidouche T, Steers NJ, Pons C, Soubeiran N, Buret D, Gilson E, Gharavi AG, D’Agati VD & Shkreli M (2024) Targeting WIP1 phosphatase promotes partial remission in experimental collapsing glomerulopathy. Kidney Int 105: 980–996

Emont MP, Jacobs C, Essene AL, Pant D, Tenen D, Colleluori G, Di Vincenzo A, Jørgensen AM, Dashti H, Stefek A, et al (2022) A single cell atlas of human and mouse white adipose tissue. Nature 603: 926–933

Ewels P, Magnusson M, Lundin S & Käller M (2016) MultiQC: summarize analysis results for multiple tools and samples in a single report. Bioinforma Oxf Engl 32: 3047–3048

Gao Z, Daquinag AC, Fussell C, Zhao Z, Dai Y, Rivera A, Snyder BE, Eckel-Mahan KL & Kolonin MG (2020) Age-associated telomere attrition in adipocyte progenitors predisposes to metabolic disease. Nat Metab 2: 1482–1497

Gao Z, Santos RB, Rupert J, Van Drunen R, Yu Y, Eckel-Mahan K & Kolonin MG (2024) Endothelial-specific telomerase inactivation causes telomere-independent cell senescence and multi-organ dysfunction characteristic of aging. Aging Cell 23: e14138

Gavia-García G, Rosado-Pérez J, Arista-Ugalde TL, Aguiñiga-Sánchez I, Santiago-Osorio E & Mendoza-Núñez VM (2021) Telomere Length and Oxidative Stress and Its Relation with Metabolic Syndrome Components in the Aging. Biology 10: 253

Ghosh A, Saginc G, Leow SC, Khattar E, Shin EM, Yan TD, Wong M, Zhang Z, Li G, Sung W-K, et al (2012) Telomerase directly regulates NF-κB-dependent transcription. Nat Cell Biol 14: 1270–1281

Hall KD & Kahan S (2018) Maintenance of lost weight and long-term management of obesity. Med Clin North Am 102: 183–197

Hao Y, Hao S, Andersen-Nissen E, Mauck WM, Zheng S, Butler A, Lee MJ, Wilk AJ, Darby C, Zager M, et al (2021) Integrated analysis of multimodal single-cell data. Cell 184: 3573–3587.e29

Hasegawa K, Zhao Y, Garbuzov A, Corces MR, Neuhöfer P, Gillespie VM, Cheung P, Belk JA, Huang Y-H, Wei Y, et al (2024) Clonal inactivation of TERT impairs stem cell competition. Nature 632: 201–208

Hernandez-Segura A, Nehme J & Demaria M (2018) Hallmarks of cellular senescence. Trends Cell Biol 28: 436–453

Hutley LJ, Newell FS, Kim Y-H, Luo X, Widberg CH, Shurety W, Prins JB & Whitehead JP (2011) A putative role for endogenous FGF-2 in FGF-1 mediated differentiation of human preadipocytes. Mol Cell Endocrinol 339: 165–171

Ikeda Y, Watanabe H, Shiuchi T, Hamano H, Horinouchi Y, Imanishi M, Goda M, Zamami Y, Takechi K, Izawa-Ishizawa Y, et al (2020) Deletion of H-ferritin in macrophages alleviates obesity and diabetes induced by high-fat diet in mice. Diabetologia 63: 1588–1602

Indran IR, Hande MP & Pervaiz S (2011) hTERT Overexpression Alleviates Intracellular ROS Production, Improves Mitochondrial Function, and Inhibits ROS-Mediated Apoptosis in Cancer Cells. Cancer Res 71: 266–276

Jacobs JJL & de Lange T (2004) Significant role for p16INK4a in p53-independent telomere-directed senescence. Curr Biol CB 14: 2302–2308

Jaitin DA, Adlung L, Thaiss CA, Weiner A, Li B, Descamps H, Lundgren P, Bleriot C, Liu Z, Deczkowska A, et al (2019) Lipid-Associated Macrophages Control Metabolic Homeostasis in a Trem2-Dependent Manner. Cell 178: 686–698.e14

Jang H, Kim M, Lee S, Kim J, Woo D-C, Kim KW, Song K & Lee I (2016) Adipose tissue hyperplasia with enhanced adipocyte-derived stem cell activity in Tc1(C8orf4)-deleted mice. Sci Rep 6: 35884

Jeffery E, Church CD, Holtrup B, Colman L & Rodeheffer MS (2015) Rapid depot-specific activation of adipocyte precursor cells at the onset of obesity. Nat Cell Biol 17: 376–385

Jeong B-C, Kim TS, Kim HS, Lee S-H & Choi Y (2015) Transmembrane protein 64 reciprocally regulates osteoblast and adipocyte differentiation by modulating Wnt/β-catenin signaling. Bone 78: 165–173

Khosravaniardakani S, Bokov DO, Mahmudiono T, Hashemi SS, Nikrad N, Rabieemotmaen S & Abbasalizad-Farhangi M (2022) Obesity Accelerates Leukocyte Telomere Length Shortening in Apparently Healthy Adults: A Meta-Analysis. Front Nutr 9: 812846

Kompella P & Vasquez KM (2019) Obesity and cancer: A mechanistic overview of metabolic changes in obesity that impact genetic instability. Mol Carcinog 58: 1531–1550

Kondoh H & Hara E (2022) Targeting p21 for diabetes: Another choice of senotherapy. Cell Metab 34: 5–7

Korsunsky I, Millard N, Fan J, Slowikowski K, Zhang F, Wei K, Baglaenko Y, Brenner M, Loh P & Raychaudhuri S (2019) Fast, sensitive and accurate integration of single-cell data with Harmony. Nat Methods 16: 1289–1296

Lagathu C, Christodoulides C, Virtue S, Cawthorn WP, Franzin C, Kimber WA, Nora ED, Campbell M, Medina-Gomez G, Cheyette BNR, et al (2009) Dact1, a nutritionally regulated preadipocyte gene, controls adipogenesis by coordinating the Wnt/beta-catenin signaling network. Diabetes 58: 609–619

Lai T-P, Zhang N, Noh J, Mender I, Tedone E, Huang E, Wright WE, Danuser G & Shay JW (2017) A method for measuring the distribution of the shortest telomeres in cells and tissues. Nat Commun 8: 1356

Lakowa N, Trieu N, Flehmig G, Lohmann T, Schön MR, Dietrich A, Zeplin PH, Langer S, Stumvoll M, Blüher M, et al (2015) Telomere length differences between subcutaneous and visceral adipose tissue in humans. Biochem Biophys Res Commun 457: 426–432

Liao X, Zhou H & Deng T (2022) The composition, function, and regulation of adipose stem and progenitor cells. J Genet Genomics 49: 308–315

Liao Y, Smyth GK & Shi W (2013) The Subread aligner: fast, accurate and scalable read mapping by seed-and-vote. Nucleic Acids Res 41: e108

Liao Y, Smyth GK & Shi W (2014) featureCounts: an efficient general purpose program for assigning sequence reads to genomic features. Bioinforma Oxf Engl 30: 923–930

Lieberman Y, Rokach L & Shay T (2018) CaSTLe - Classification of single cells by transfer learning: Harnessing the power of publicly available single cell RNA sequencing experiments to annotate new experiments. PloS One 13: e0205499

Lin S, Nascimento EM, Gajera CR, Chen L, Neuhöfer P, Garbuzov A, Wang S & Artandi SE (2018) Distributed hepatocytes expressing telomerase repopulate the liver in homeostasis and injury. Nature 556: 244–248

Lipskaia L, Breau M, Cayrou C, Churikov D, Braud L, Jacquet J, Born E, Fouillade C, Curras-Alonso S, Bauwens S, et al (2024) mTert induction in p21-positive cells counteracts capillary rarefaction and pulmonary emphysema. EMBO Rep: 1–35

Liu TM, Lee EH, Lim B & Shyh-Chang N (2016) Concise Review: Balancing Stem Cell Self-Renewal and Differentiation with PLZF. Stem Cells Dayt Ohio 34: 277–287

Liu Z, Wu KKL, Jiang X, Xu A & Cheng KKY (2020) The role of adipose tissue senescence in obesity- and ageing-related metabolic disorders. Clin Sci 134: 315–330

Longo M, Zatterale F, Naderi J, Parrillo L, Formisano P, Raciti GA, Beguinot F & Miele C (2019) Adipose Tissue Dysfunction as Determinant of Obesity-Associated Metabolic Complications. Int J Mol Sci 20: 2358

López-Otín C, Blasco MA, Partridge L, Serrano M & Kroemer G (2013) The Hallmarks of Aging. Cell 153: 1194–1217

Love MI, Huber W & Anders S (2014) Moderated estimation of fold change and dispersion for RNA-seq data with DESeq2. Genome Biol 15: 550

Lu B, Guo S, Zhao J, Wang X & Zhou B (2024) Adipose knockout of H-ferritin improves energy metabolism in mice. Mol Metab 80: 101871

Lun ATL, Riesenfeld S, Andrews T, Dao TP, Gomes T, participants in the 1st Human Cell Atlas Jamboree & Marioni JC (2019) EmptyDrops: distinguishing cells from empty droplets in droplet-based single-cell RNA sequencing data. Genome Biol 20: 63

Ma F & Pellegrini M (2020) ACTINN: automated identification of cell types in single cell RNA sequencing. Bioinforma Oxf Engl 36: 533–538

Magalhaes MS, Smith P, Portman JR, Jackson-Jones LH, Bain CC, Ramachandran P, Michailidou Z, Stimson RH, Dweck MR, Denby L, et al (2021) Role of Tim4 in the regulation of ABCA1+ adipose tissue macrophages and post-prandial cholesterol levels. Nat Commun 12: 4434

Maniyadath B, Zhang Q, Gupta RK & Mandrup S (2023) Adipose tissue at single-cell resolution. Cell Metab 35: 386–413

Marcelin G & Clément K (2021) The multifaceted progenitor fates in healthy or unhealthy adipose tissue during obesity. Rev Endocr Metab Disord 22: 1111–1119

McCarthy DJ, Campbell KR, Lun ATL & Wills QF (2017) Scater: pre-processing, quality control, normalization and visualization of single-cell RNA-seq data in R. Bioinforma Oxf Engl 33: 1179–1186

McGinnis CS, Murrow LM & Gartner ZJ (2019) DoubletFinder: Doublet Detection in Single-Cell RNA Sequencing Data Using Artificial Nearest Neighbors. Cell Syst 8: 329–337.e4

McNelis JC & Olefsky JM (2014) Macrophages, Immunity, and Metabolic Disease. Immunity 41: 36–48

Merrick D, Sakers A, Irgebay Z, Okada C, Calvert C, Morley MP, Percec I & Seale P (2019) Identification of a mesenchymal progenitor cell hierarchy in adipose tissue. Science 364: eaav2501

Mirzaeicheshmeh E, Zerrweck C, Centeno-Cruz F, Baca-Peynado P, Martinez-Hernandez A, García-Ortiz H, Contreras-Cubas C, Salas-Martínez MG, Saldaña-Alvarez Y, Mendoza-Caamal EC, et al (2021) Alterations of DNA methylation during adipogenesis differentiation of mesenchymal stem cells isolated from adipose tissue of patients with obesity is associated with type 2 diabetes. Adipocyte 10: 493–504

Montandon M, Hamidouche T, Yart L, Duret LC, Pons C, Soubeiran N, Pousse M, Cervera L, Vial V, Fassy J, et al (2022) Telomerase is required for glomerular renewal in kidneys of adult mice. NPJ Regen Med 7: 15

de Morree A & Rando TA (2023) Regulation of adult stem cell quiescence and its functions in the maintenance of tissue integrity. Nat Rev Mol Cell Biol 24: 334–354

Murray PJ (2017) Macrophage Polarization. Annu Rev Physiol 79: 541–566

Nahta R & Castellino RC (2021) Phosphatase magnesium-dependent 1 δ (PPM1D), serine/threonine protein phosphatase and novel pharmacological target in cancer. Biochem Pharmacol 184: 114362

Narasimhan A, Flores RR, Camell CD, Bernlohr DA, Robbins PD & Niedernhofer LJ (2022) Cellular Senescence in Obesity and Associated Complications: a New Therapeutic Target. Curr Diab Rep 22: 537– 548

Nee K, Nguyen Q & Kessenbrock K (2018) Single Nuclei RNA Sequencing of Breast Adipose Tissue (10x Nuclei-Seq).

Neuhöfer P, Roake CM, Kim SJ, Lu RJ, West RB, Charville GW & Artandi SE (2021) Acinar cell clonal expansion in pancreas homeostasis and carcinogenesis. Nature 597: 715–719

Palani NP, Horvath C, Timshel PN, Folkertsma P, Grønning AGB, Henriksen TI, Peijs L, Jensen VH, Sun W, Jespersen NZ, et al (2023) Adipogenic and SWAT cells separate from a common progenitor in human brown and white adipose depots. Nat Metab 5: 996–1013

Palhinha L, Liechocki S, Hottz ED, Pereira JA da S, de Almeida CJ, Moraes-Vieira PMM, Bozza PT & Maya-Monteiro CM (2019) Leptin Induces Proadipogenic and Proinflammatory Signaling in Adipocytes. Front Endocrinol 10: 841

Palmer AK, Tchkonia T, LeBrasseur NK, Chini EN, Xu M & Kirkland JL (2015) Cellular Senescence in Type 2 Diabetes: A Therapeutic Opportunity. Diabetes 64: 2289–2298

Palmer AK, Xu M, Zhu Y, Pirtskhalava T, Weivoda MM, Hachfeld CM, Prata LG, van Dijk TH, Verkade E, Casaclang-Verzosa G, et al (2019) Targeting senescent cells alleviates obesity-induced metabolic dysfunction. Aging Cell 18

Park J-I, Venteicher AS, Hong JY, Choi J, Jun S, Shkreli M, Chang W, Meng Z, Cheung P, Ji H, et al (2009) Telomerase modulates Wnt signalling by association with target gene chromatin. Nature 460: 66–72

Pettersson US, Waldén TB, Carlsson P-O, Jansson L & Phillipson M (2012) Female Mice are Protected against High-Fat Diet Induced Metabolic Syndrome and Increase the Regulatory T Cell Population in Adipose Tissue. PLoS ONE 7: e46057

Pini M, Czibik G, Sawaki D, Mezdari Z, Braud L, Delmont T, Mercedes R, Martel C, Buron N, Marcelin G, et al (2021) Adipose tissue senescence is mediated by increased ATP content after a short-term high-fat diet exposure. Aging Cell 20: e13421

Polański K, Young MD, Miao Z, Meyer KB, Teichmann SA & Park J-E (2020) BBKNN: fast batch alignment of single cell transcriptomes. Bioinformatics 36: 964–965

Riant E, Waget A, Cogo H, Arnal J-F, Burcelin R & Gourdy P (2009) Estrogens Protect against High-Fat Diet-Induced Insulin Resistance and Glucose Intolerance in Mice. Endocrinology 150: 2109–2117

Ronti T, Lupattelli G & Mannarino E (2006) The endocrine function of adipose tissue: an update. Clin Endocrinol (Oxf) 64: 355–365

Sadler AJ, Rossello FJ, Yu L, Deane JA, Yuan X, Wang D, Irving AT, Kaparakis-Liaskos M, Gantier MP, Ying H, et al (2015) BTB-ZF transcriptional regulator PLZF modifies chromatin to restrain inflammatory signaling programs. Proc Natl Acad Sci 112: 1535–1540

Sahin E, Colla S, Liesa M, Moslehi J, Müller FL, Guo M, Cooper M, Kotton D, Fabian AJ, Walkey C, et al (2011) Telomere dysfunction induces metabolic and mitochondrial compromise. Nature 470: 359–365

Sakers A, De Siqueira MK, Seale P & Villanueva CJ (2022) Adipose-tissue plasticity in health and disease. Cell 185: 419–446

Saretzki G (2014) Extra-telomeric functions of human telomerase: cancer, mitochondria and oxidative stress. Curr Pharm Des 20: 6386–6403

Sarin KY, Cheung P, Gilison D, Lee E, Tennen RI, Wang E, Artandi MK, Oro AE & Artandi SE (2005) Conditional telomerase induction causes proliferation of hair follicle stem cells. Nature 436: 1048–1052

Sárvári AK, Van Hauwaert EL, Markussen LK, Gammelmark E, Marcher A-B, Ebbesen MF, Nielsen R, Brewer JR, Madsen JGS & Mandrup S (2021) Plasticity of Epididymal Adipose Tissue in Response to Diet-Induced Obesity at Single-Nucleus Resolution. Cell Metab 33: 437–453.e5

Schafer MJ, White TA, Evans G, Tonne JM, Verzosa GC, Stout MB, Mazula DL, Palmer AK, Baker DJ, Jensen MD, et al (2016) Exercise Prevents Diet-Induced Cellular Senescence in Adipose Tissue. Diabetes 65: 1606–1615

Ségal-Bendirdjian E & Geli V (2019) Non-canonical Roles of Telomerase: Unraveling the Imbroglio. Front Cell Dev Biol 7

Shanmugam R, Majee P, Shi W, Ozturk MB, Vaiyapuri TS, Idzham K, Raju A, Shin SH, Fidan K, Low J-L, et al (2024) Iron-(Fe3+) dependent reactivation of telomerase drives colorectal cancers. Cancer Discov

Shao M, Vishvanath L, Busbuso NC, Hepler C, Shan B, Sharma AX, Chen S, Yu X, An YA, Zhu Y, et al (2018) De novo adipocyte differentiation from Pdgfrβ+ preadipocytes protects against pathologic visceral adipose expansion in obesity. Nat Commun 9: 890

Shim HS, Iaconelli J, Shang X, Li J, Lan ZD, Jiang S, Nutsch K, Beyer BA, Lairson LL, Boutin AT, et al (2024) TERT activation targets DNA methylation and multiple aging hallmarks. Cell

Shkreli M, Sarin KY, Pech MF, Papeta N, Chang W, Brockman SA, Cheung P, Lee E, Kuhnert F, Olson JL, et al (2011) Reversible cell cycle entry in adult kidney podocytes through regulated control of telomerase and Wnt signaling. Nat Med 18: 111–119

Shreeram S, Demidov ON, Hee WK, Yamaguchi H, Onishi N, Kek C, Timofeev ON, Dudgeon C, Fornace AJ, Anderson CW, et al (2006) Wip1 phosphatase modulates ATM-dependent signaling pathways. Mol Cell 23: 757–764

Spinelli R, Baboota RK, Gogg S, Beguinot F, Blüher M, Nerstedt A & Smith U Increased cell senescence in human metabolic disorders. J Clin Invest 133: e169922

Sturmlechner I, Zhang C, Sine CC, van Deursen E-J, Jeganathan KB, Hamada N, Grasic J, Friedman D, Stutchman JT, Can I, et al (2021) p21 produces a bioactive secretome that places stressed cells under immunosurveillance. Science 374: eabb3420

Subramanian A, Tamayo P, Mootha VK, Mukherjee S, Ebert BL, Gillette MA, Paulovich A, Pomeroy SL, Golub TR, Lander ES, et al (2005) Gene set enrichment analysis: a knowledge-based approach for interpreting genome-wide expression profiles. Proc Natl Acad Sci U S A 102: 15545–15550

Sun K, Kusminski CM & Scherer PE (2011) Adipose tissue remodeling and obesity. J Clin Invest 121: 2094– 2101

Sun L, Chiang JY, Choi JY, Xiong Z-M, Mao X, Collins FS, Hodes RJ & Cao K (2019) Transient induction of telomerase expression mediates senescence and reduces tumorigenesis in primary fibroblasts. Proc Natl Acad Sci U S A 116: 18983–18993

Tabula Muris Consortium, Overall coordination, Logistical coordination, Organ collection and processing, Library preparation and sequencing, Computational data analysis, Cell type annotation, Writing group, Supplemental text writing group, & Principal investigators (2018) Single-cell transcriptomics of 20 mouse organs creates a Tabula Muris. Nature 562: 367–372

van Tienen FHJ, Laeremans H, van der Kallen CJH & Smeets HJM (2009) Wnt5b stimulates adipogenesis by activating PPARgamma, and inhibiting the beta-catenin dependent Wnt signaling pathway together with Wnt5a. Biochem Biophys Res Commun 387: 207–211

Vergoni B, Cornejo P-J, Gilleron J, Djedaini M, Ceppo F, Jacquel A, Bouget G, Ginet C, Gonzalez T, Maillet J, et al (2016) DNA Damage and the Activation of the p53 Pathway Mediate Alterations in Metabolic and Secretory Functions of Adipocytes. Diabetes 65: 3062–3074

Vishvanath L & Gupta RK (2019) Contribution of adipogenesis to healthy adipose tissue expansion in obesity. J Clin Invest 129: 4022–4031

Wang L, Wang B, Gasek NS, Zhou Y, Cohn RL, Martin DE, Zuo W, Flynn WF, Guo C, Jellison ER, et al (2021) Targeting p21Cip1 highly expressing cells in adipose tissue alleviates insulin resistance in obesity. Cell Metab

Warnes GR, Bolker B, Bonebakker L, Gentleman R, Huber W, Liaw A, Lumley T, Maechler M, Magnusson A & Moeller S (2009) gplots: Various R programming tools for plotting data. R Package Version 2: 1

Weisberg SP, McCann D, Desai M, Rosenbaum M, Leibel RL & Ferrante AW (2003) Obesity is associated with macrophage accumulation in adipose tissue. J Clin Invest 112: 1796–1808

Wu T, Hu E, Xu S, Chen M, Guo P, Dai Z, Feng T, Zhou L, Tang W, Zhan L, et al (2021) clusterProfiler 4.0: A universal enrichment tool for interpreting omics data. Innov Camb Mass 2: 100141

Zhu L, Brown WC, Cai Q, Krust A, Chambon P, McGuinness OP & Stafford JM (2013) Estrogen Treatment After Ovariectomy Protects Against Fatty Liver and May Improve Pathway-Selective Insulin Resistance. Diabetes 62: 424–434

