## Supplementary figures and images for "TERT prevents obesity-induced metabolic disorders by promoting adipose stem cell expansion and differentiation"

### Suppl Figure 1

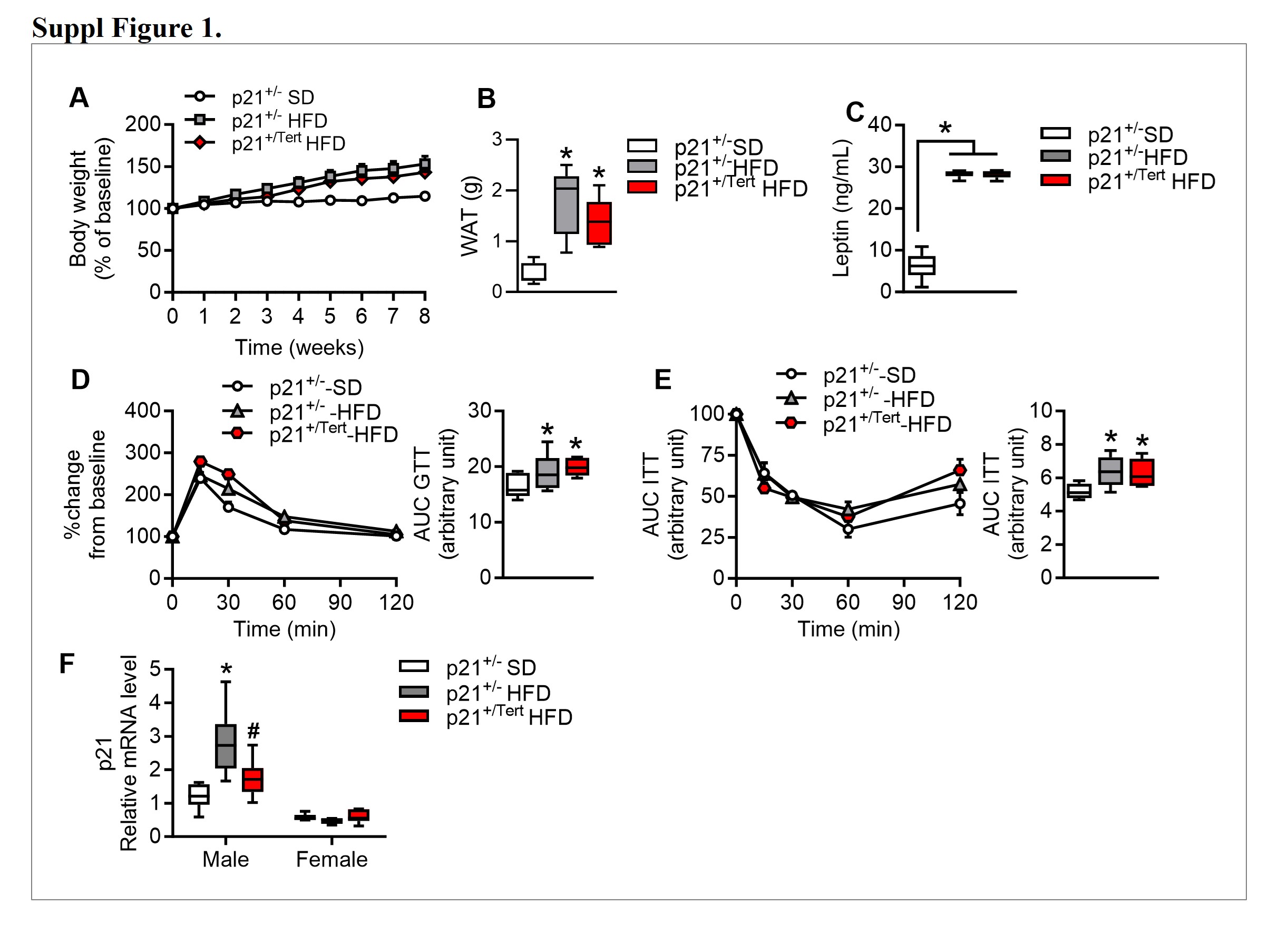

### Suppl Figure 2

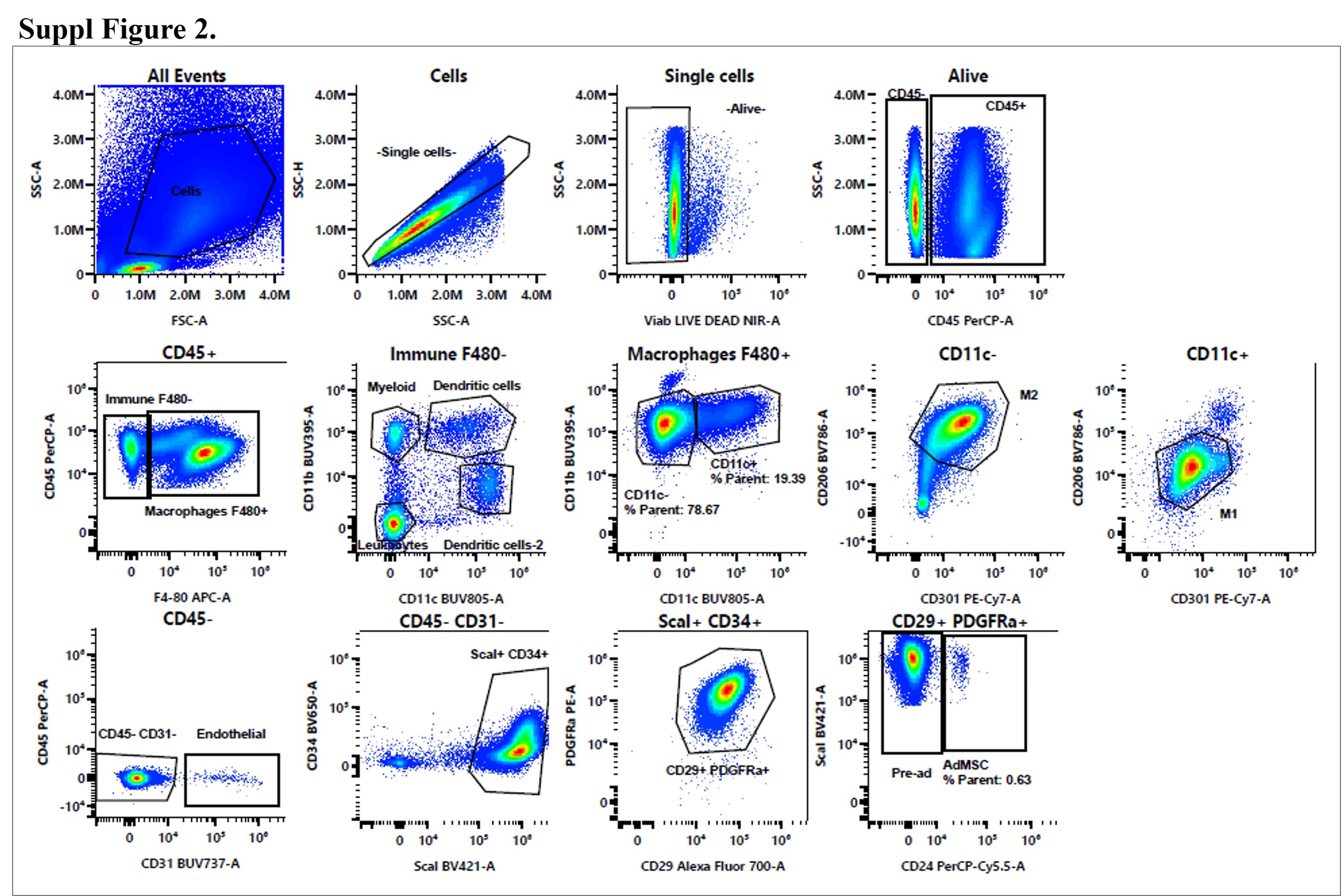

### Suppl Figure 3

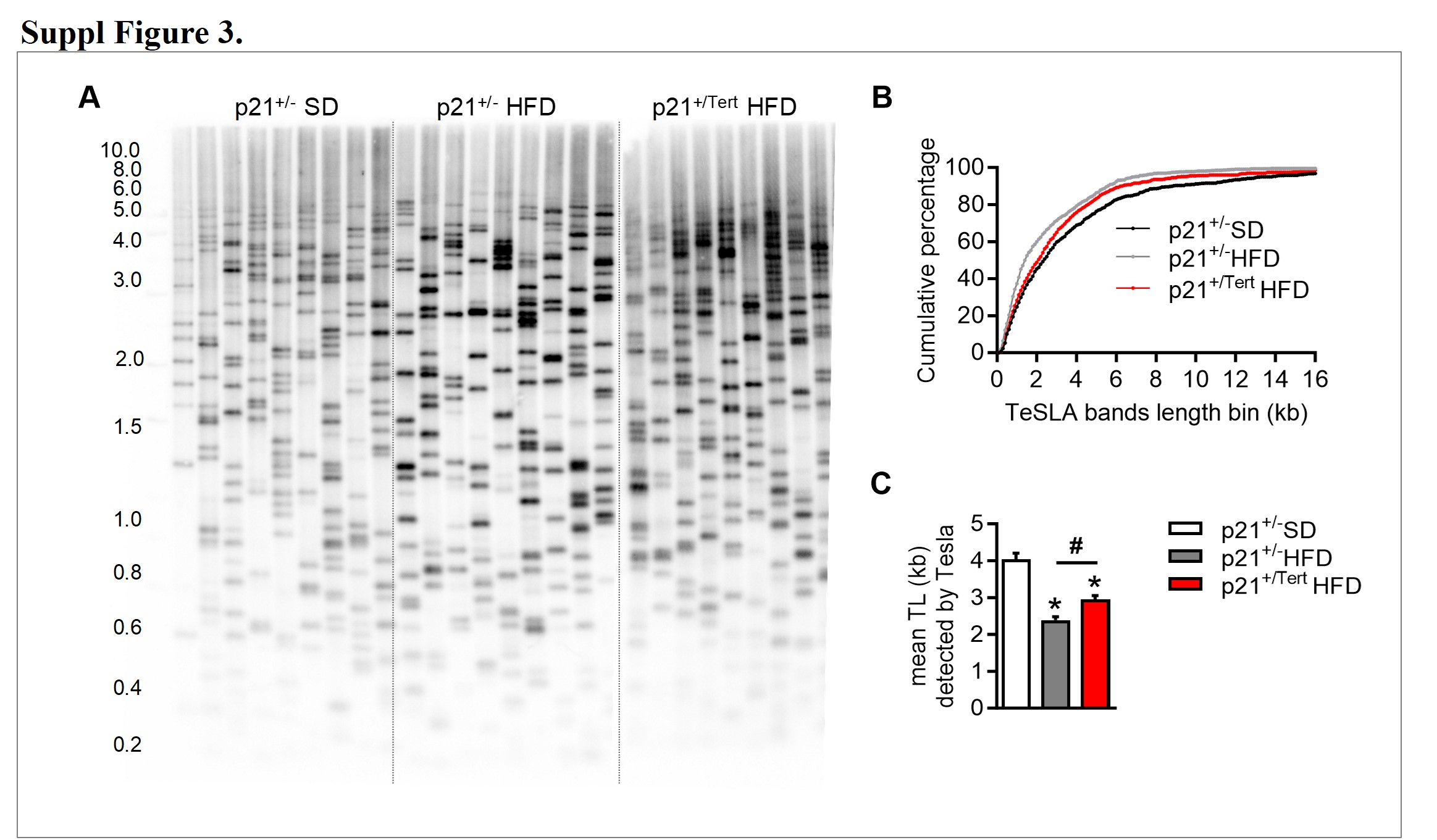

### Suppl Figure 4

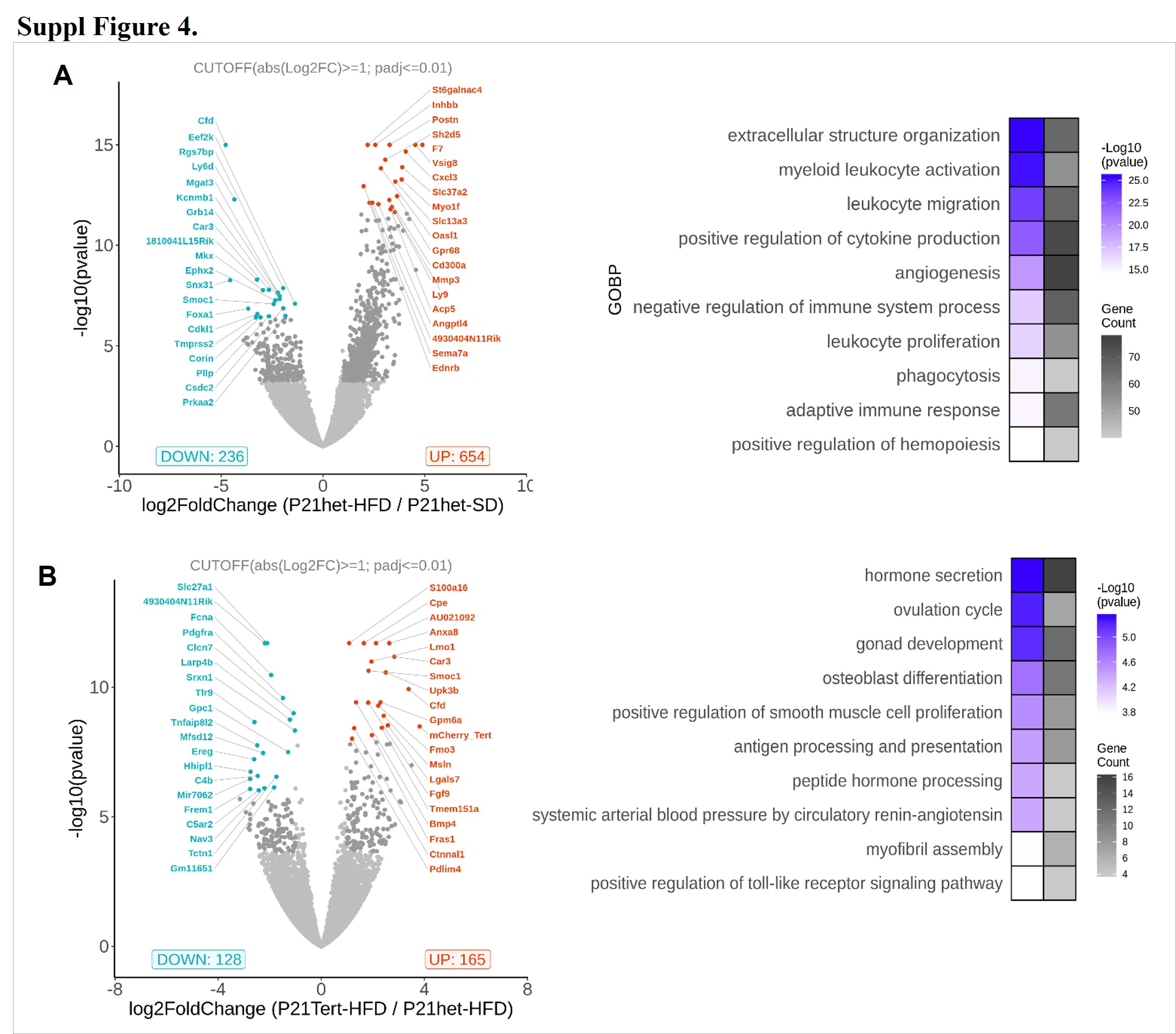

### Suppl Figure 5

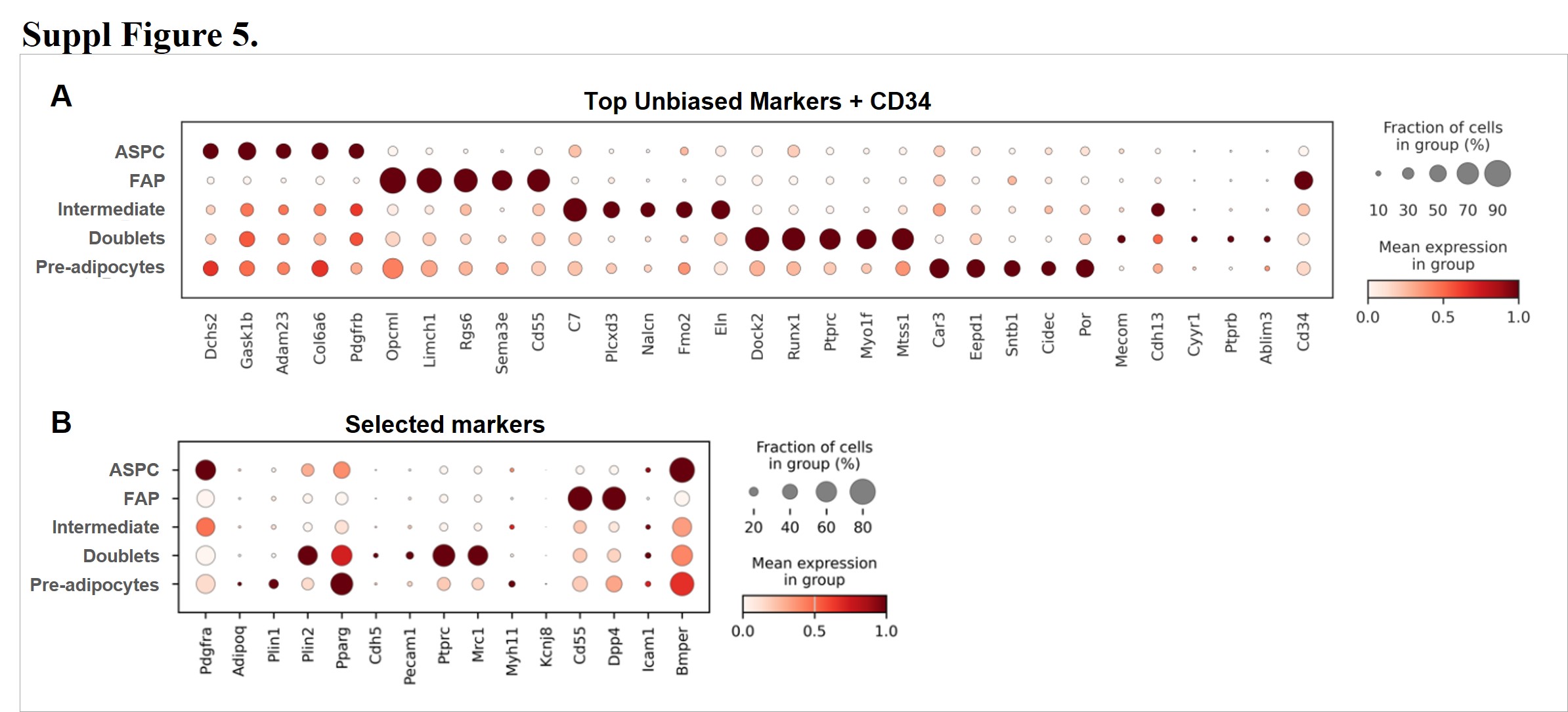

### Suppl Figure 6

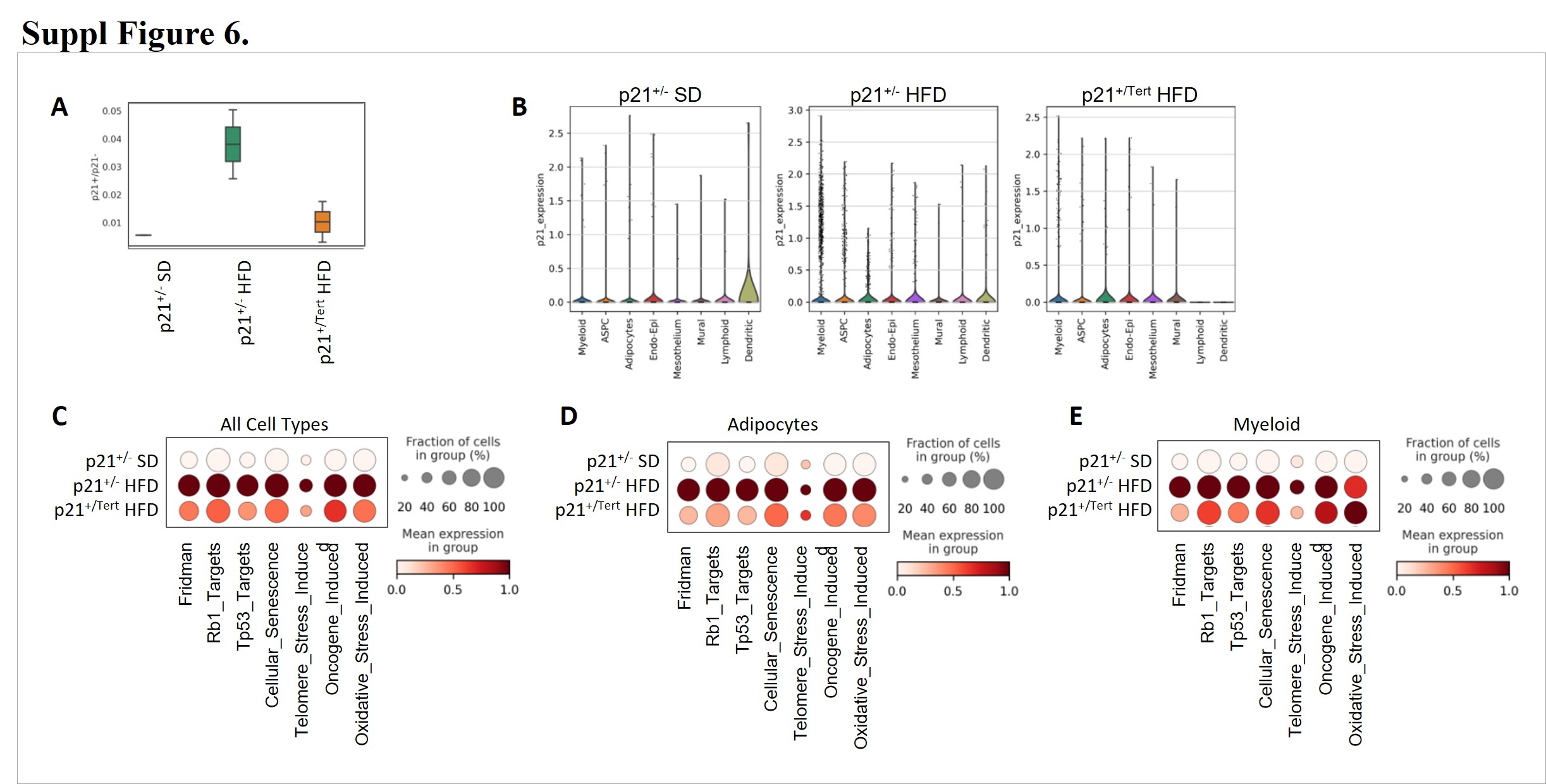

### Suppl Figure 7

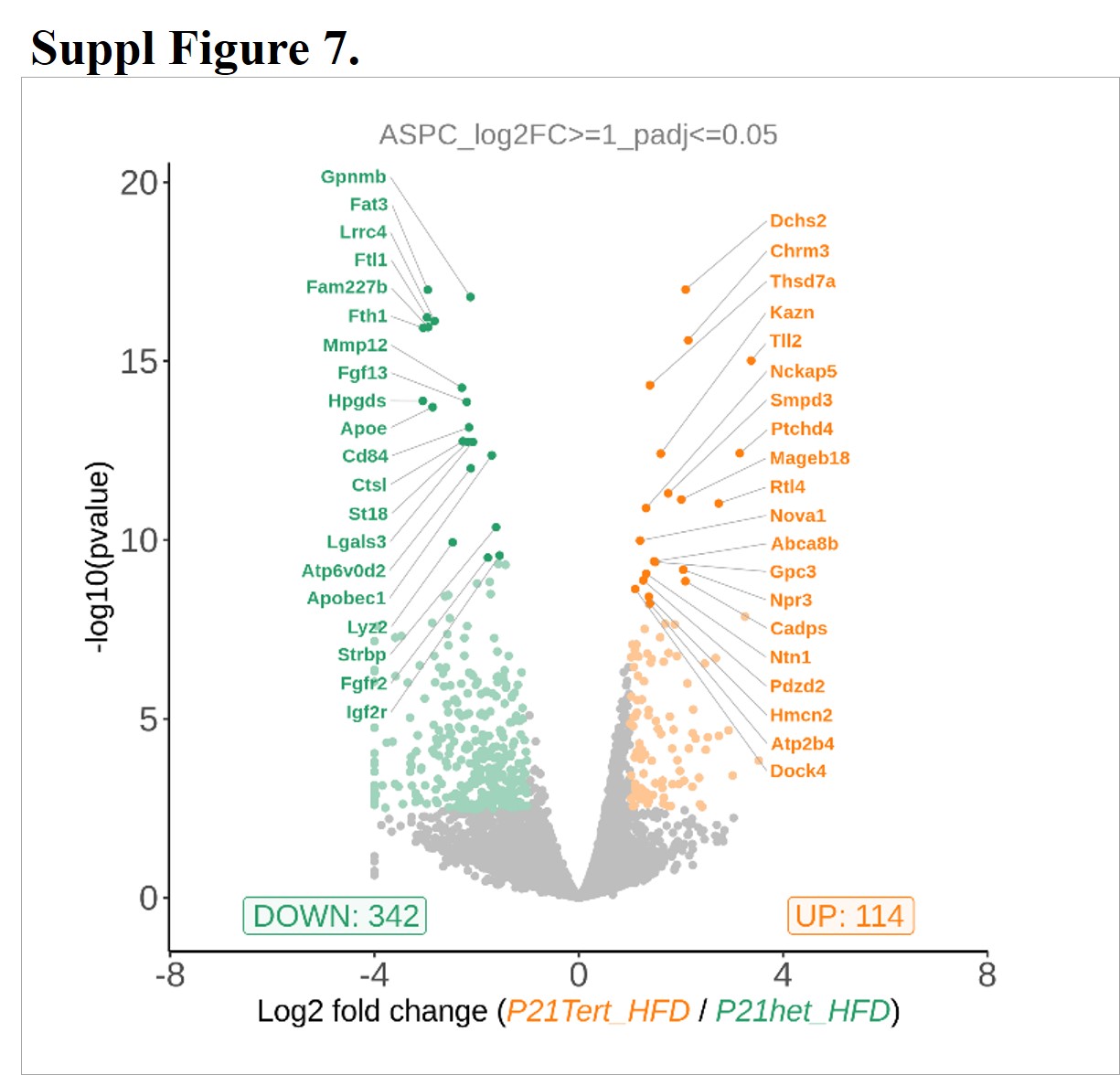

### Suppl Figure 8

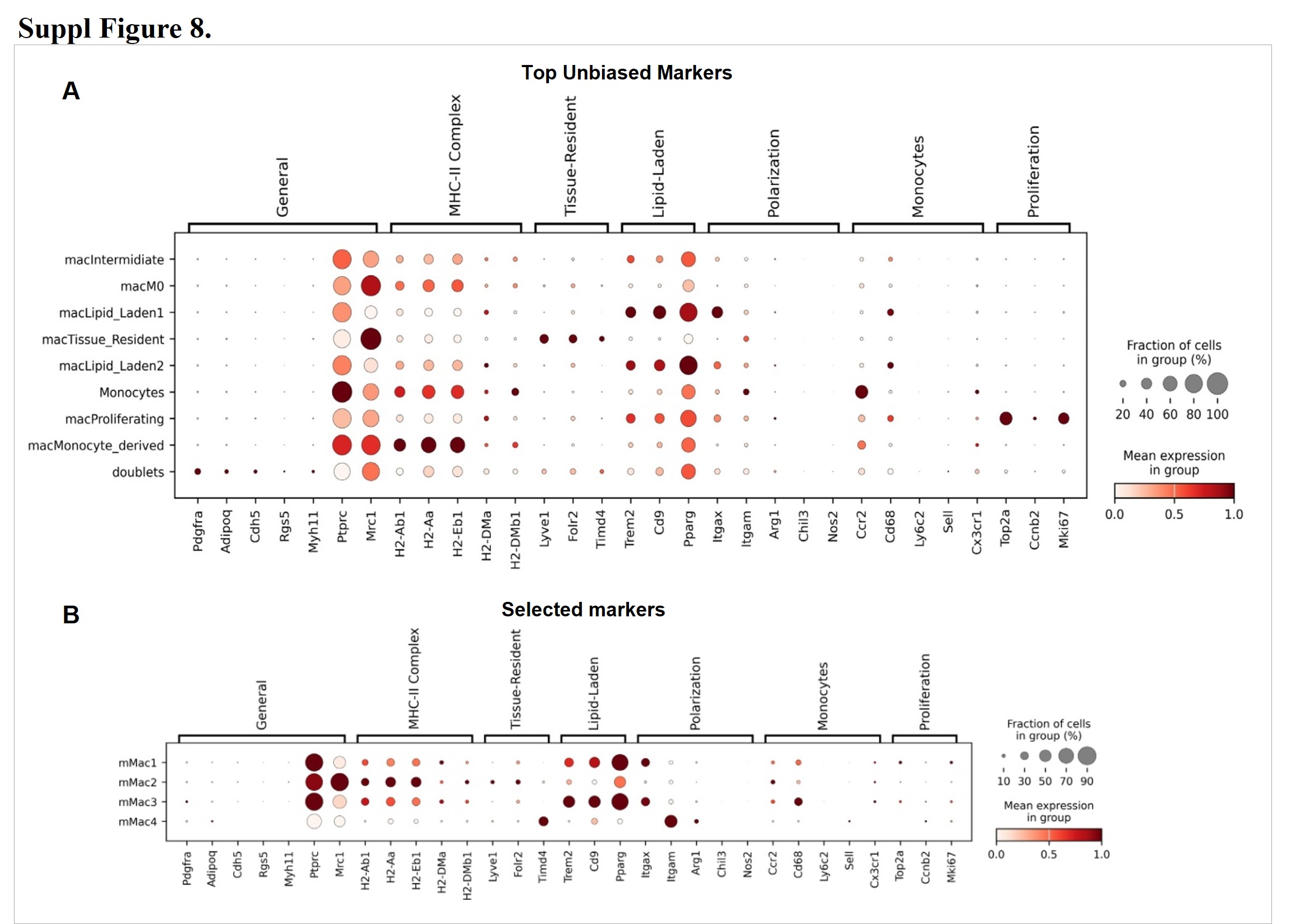

### Suppl Figure 9

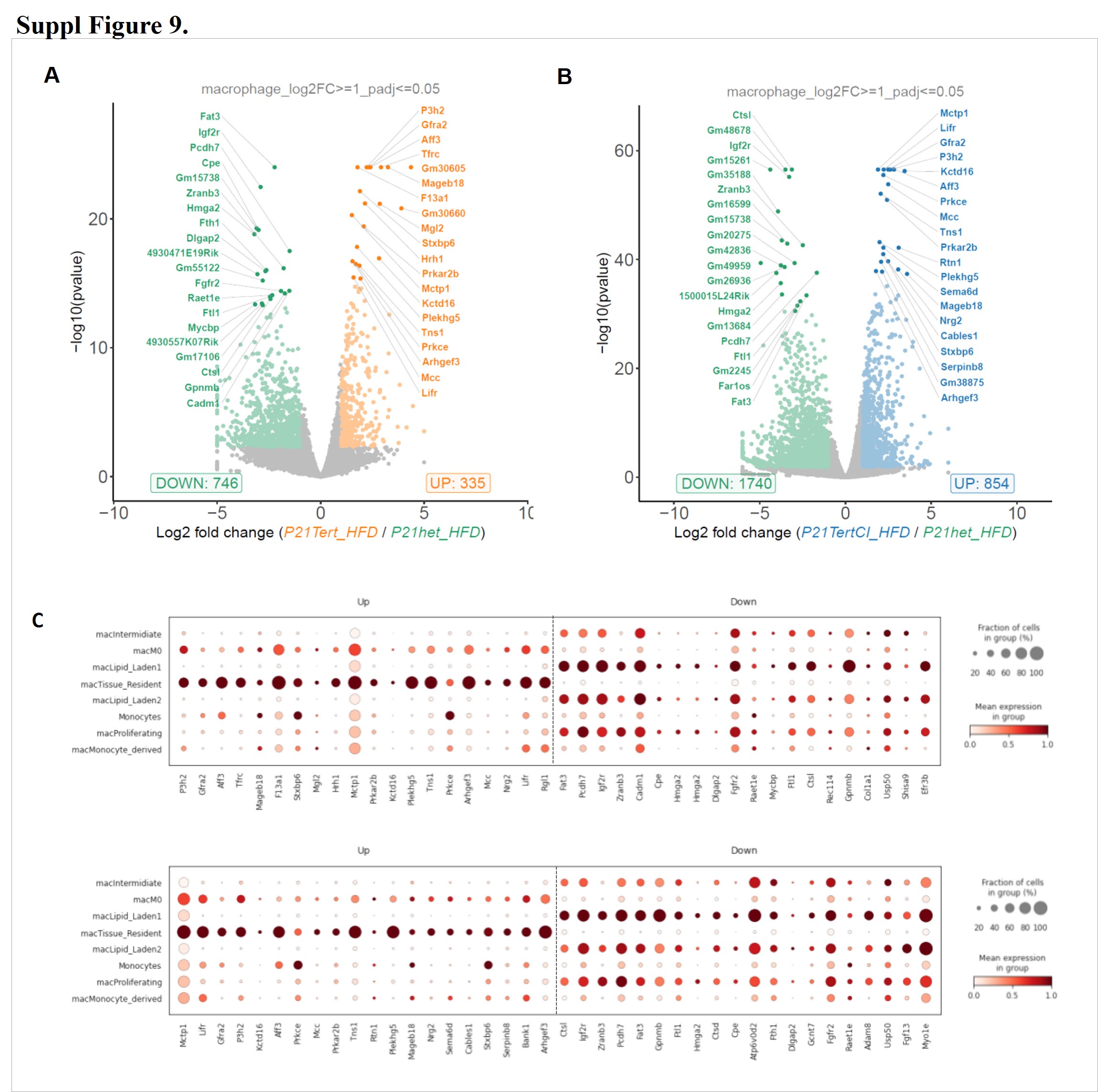

### Suppl Figure 10

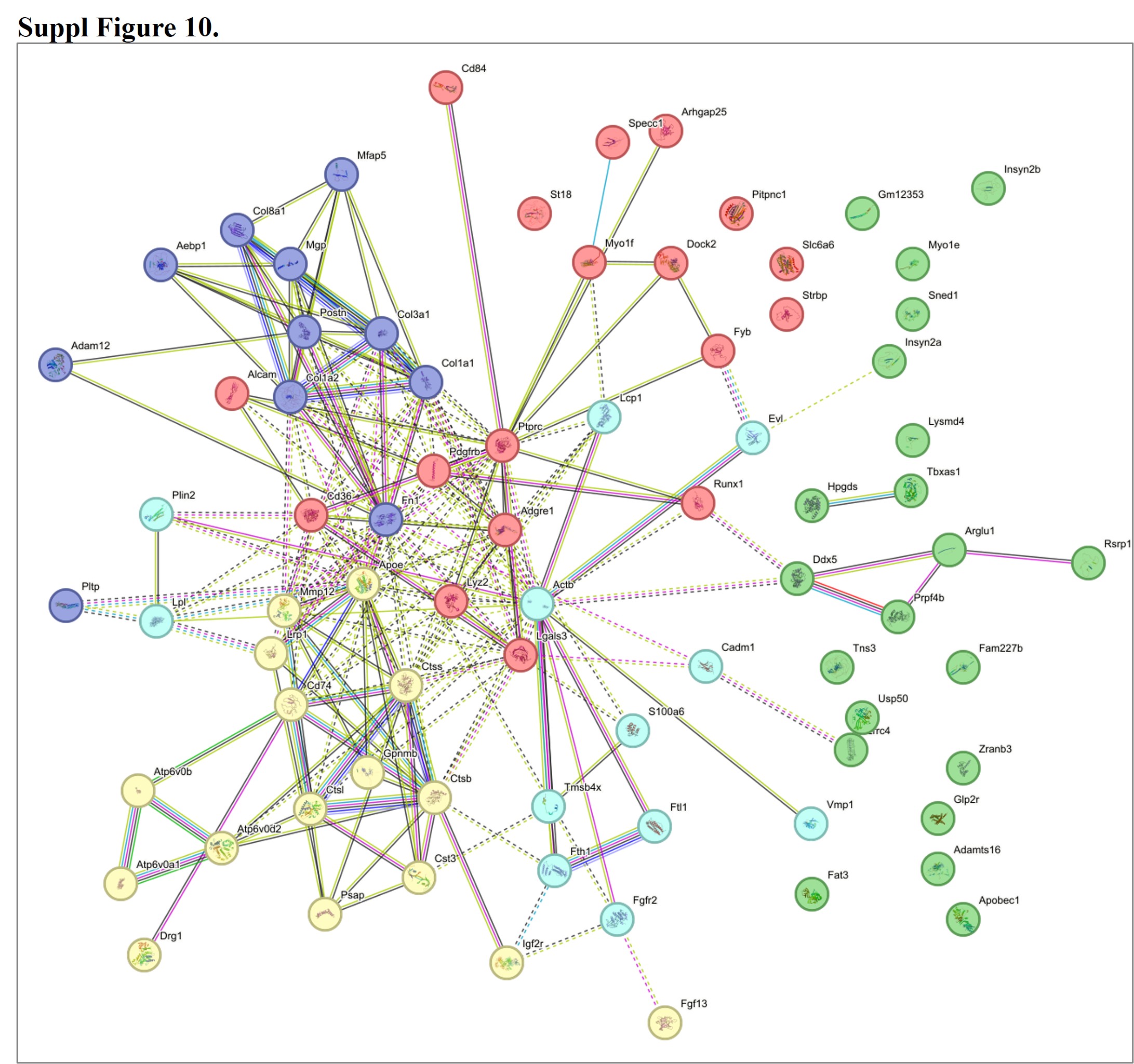

### Suppl Figure 11

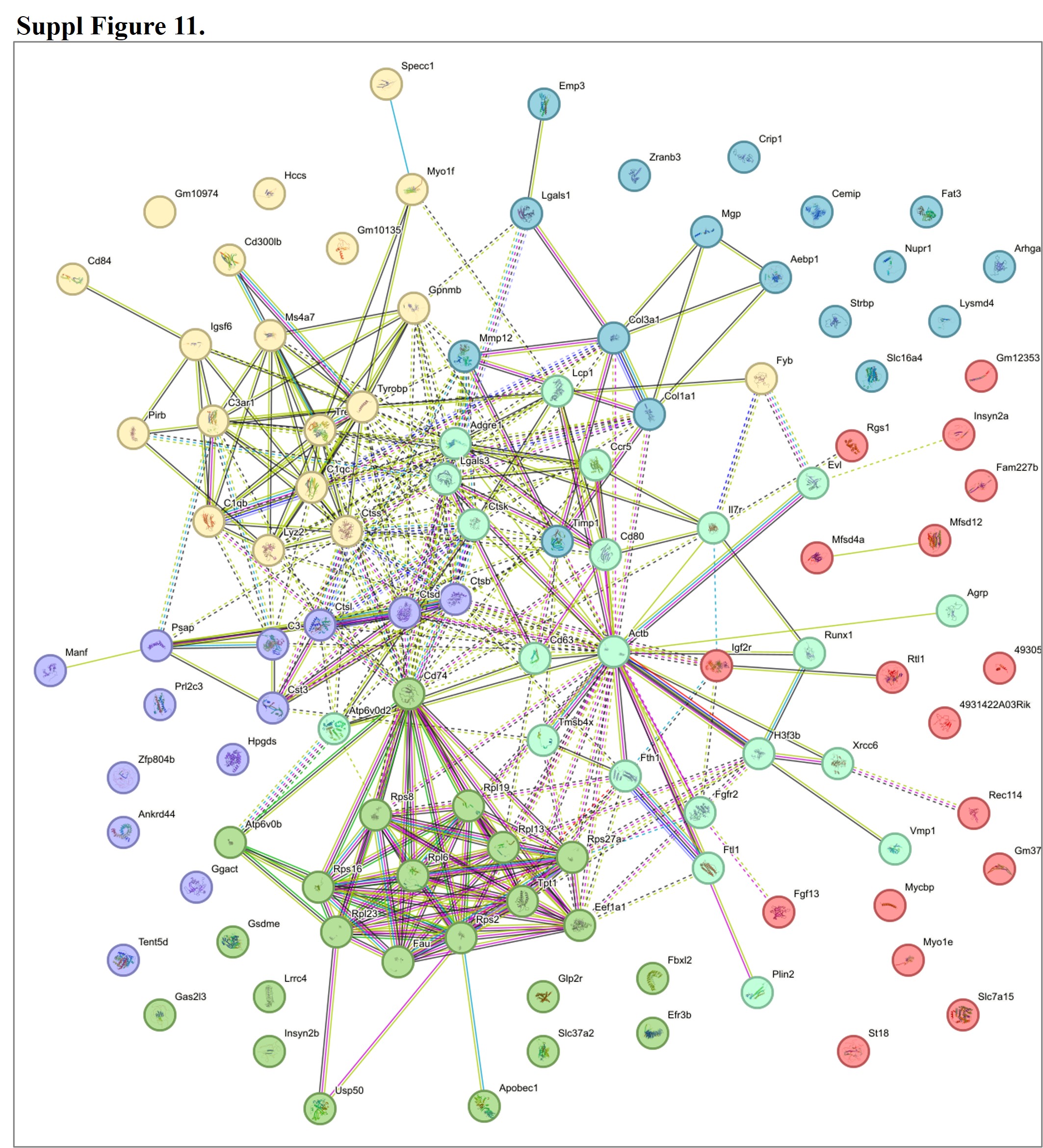
